# SALTS – <u>S</u>URFR (sncRNA) <u>A</u>nd <u>L</u>AGOOn (lncRNA) <u>T</u>ranscriptomics <u>S</u>uite

**DOI:** 10.1101/2021.02.08.430280

**Authors:** Mohan V Kasukurthi, Dominika Houserova, Yulong Huang, Addison A. Barchie, Justin T. Roberts, Dongqi Li, Bin Wu, Jingshan Huang, Glen M Borchert

## Abstract

The widespread utilization of high-throughput sequencing technologies has unequivocally demonstrated that eukaryotic transcriptomes consist primarily (>98%) of non-coding RNA (ncRNA) transcripts significantly more diverse than their protein-coding counterparts.

ncRNAs are typically divided into two categories based on their length. (1) ncRNAs less than 200 nucleotides (nt) long are referred as small non-coding RNAs (sncRNAs) and include microRNAs (miRNAs), piwi-interacting RNAs (piRNAs), small nucleolar RNAs (snoRNAs), transfer ribonucleic RNAs (tRNAs), etc., and the majority of these are thought to function primarily in controlling gene expression. That said, the full repertoire of sncRNAs remains fairly poorly defined as evidenced by two entirely new classes of sncRNAs only recently being reported, i.e., snoRNA-derived RNAs (sdRNAs) and tRNA-derived fragments (tRFs). (2) ncRNAs longer than 200 nt long are known as long ncRNAs (lncRNAs). lncRNAs represent the 2^nd^ largest transcriptional output of the cell (behind only ribosomal RNAs), and although functional roles for several lncRNAs have been reported, most lncRNAs remain largely uncharacterized due to a lack of predictive tools aimed at guiding functional characterizations.

Importantly, whereas the cost of high-throughput transcriptome sequencing is now feasible for most active research programs, tools necessary for the interpretation of these sequencings typically require significant computational expertise and resources markedly hindering widespread utilization of these datasets. In light of this, we have developed a powerful new ncRNA transcriptomics suite, SALTS, which is highly accurate, markedly efficient, and extremely user-friendly. SALTS stands for <u>S</u>URFR (sncRNA) <u>A</u>nd <u>L</u>AGOOn (lncRNA) <u>T</u>ranscriptomics <u>S</u>uite and offers platforms for comprehensive sncRNA and lncRNA profiling and discovery, ncRNA functional prediction, and the identification of significant differential expressions among datasets. Notably, SALTS is accessed through an intuitive Web-based interface, can be used to analyze either user-generated, standard next-generation sequencing (NGS) output file uploads (e.g., FASTQ) or existing NCBI Sequence Read Archive (SRA) data, and requires absolutely no dataset pre-processing or knowledge of library adapters/oligonucleotides.

SALTS constitutes the first publically available, Web-based, comprehensive ncRNA transcriptomic NGS analysis platform designed specifically for users with no computational background, providing a much needed, powerful new resource capable of enabling more widespread ncRNA transcriptomic analyses. The SALTS WebServer is freely available online at http://salts.soc.southalabama.edu.

## GENERAL INTRODUCTION

Cellular metabolism and survival are greatly dependent on how quickly and efficiently the cell can respond to internal and external stimuli. This process often requires tightly orchestrated genome-wide changes in gene expression. With rapid technological advancements in both genomics and transcriptomics, particularly the development of robust deep sequencing, it is ever more apparent that many regulatory non-coding RNAs (ncRNAs) that help coordinate gene expression changes remain elusive and the networks created thereof are far more complex than previously thought(1). As many of these are dynamic and their presence or absence is highly conditional (i.e., environmental stress, disease, tissue type, etc.), their identification poses a challenge and many remain undescribed(2). As such, we have developed a set of guidelines and parameters to help confidently identify and characterize these molecules. Importantly, by implementing alternative strategies for next-generation sequencing (NGS) analysis based on examining conditional changes in expression and/or fragmentation patterns from individual genomic loci rather than depending on pre-existing annotations, we find previously elusive ncRNAs can now be readily identified via our platform. In addition to this, we have also developed an array of downstream analyses to more fully characterize identified ncRNAs and predict their functional roles (e.g., molecular targets).

To date several platforms aimed at either small non-coding RNA (sncRNA) or long non-coding RNA (lncRNA) characterization have been developed(3). Although each of these existing platforms possess some unique advantages, each also carry their own critical limitations (detailed herein). That said, to our knowledge, SALTS is the first-ever resource designed to determine ncRNA expressions in both short ncRNA-Seq and standard RNA-Seq datasets and to provide functional predictions for ncRNAs identified in either. Perhaps most importantly, however, in addition to being highly accurate and efficient, SALTS has been developed to require absolutely no computational background in order to enable widespread ncRNA transcriptomic analysis by a much broader community of researchers. Of note, a clear, step-by-step user manual for the SALTS platform is provided in **Supplemental Information File 1**.

### SECTION 1. SALTS Tool for Small non-coding RNA Analysis: SURFR

ncRNAs less than 200 nucleotides (nt) in length are referred to as small non-coding RNAs (sncRNAs) and include microRNAs (miRNAs), piwi-interacting RNAs (piRNAs), small nucleolar RNAs (snoRNAs), transfer ribonucleic RNAs (tRNAs), etc.(4). One striking example of the regulatory capabilities of sncRNAs comes from a group of small yet potent RNAs called microRNAs (miRNAs). MiRNAs are ~22 nt RNAs excised from longer pre-miRNA hairpins that function through associating with the RNA-induced silencing complex (RISC) in order to bind to the 3’ UTRs of their target mRNAs and repress their translational activities(5). In just the past two decades, thousands of miRNAs have been identified and implicated in regulating cell growth, differentiation, and apoptosis(6), as well as contributing to tumorigenesis(7) and chemoresistance(8). As this group has been thoroughly examined due to its relevance to various types of cancer(9), it is now widely accepted that a single miRNA is capable of altering the expression of whole cohorts of protein coding genes(4). Importantly, studies aimed at evaluating the transcriptomic changes of miRNAs have revealed the existence of miRNA-like fragments derived from other ncRNA biotypes and suggest similar regulatory capacities may be associated with these novel sncRNAs(10–13). As such, we suggest that the SURFR resource described herein represents an intuitive, high throughput platform capable of revisiting old NGS datasets and identifying novel, relevant miRNA-like fragments derived from other types of ncRNAs that were previously overlooked.

Comparably sized, miRNA-like fragments excised from many other types of ncRNAs have now been reported and many of these shown to similarly regulate gene expressions and/or chromatin compaction (e.g., piRNAs, rasiRNAs, rRNAs, scRNAs, snoRNAs, snRNAs, RNase P, tRNAs, Y RNAs, and Vault RNAs)(10–13). That said, the expressions and functions of the vast majority of specific sncRNA fragments excised from anything other than annotated miRNAs remain largely undefined, although fragments from snoRNAs (sdRNAs) and tRNAs (tRFs) have recently begun to receive considerably more attention(12, 13). In 2008, Ender et al. were the first to report a small RNA fragment originating from a snoRNA, ACA45(14). Despite the principle snoRNA function being long characterized as guiding rRNA modifications, they showed that this snoRNA-derived RNA (sdRNA) was not only processed by Dicer-like regular miRNAs but also capable of silencing CDC2L6 gene in miRNA-like manner. Since then various other studies have described similar fragments arising from other snoRNAs (reviewed in (15)) as well as from other types of ncRNAs. Notably, tRNA-derived fragments (tRFs) have recently gained attention due to their differential abundance under highly specific conditions, such as developmental stage(16), stress(17), or viral infection(18). Moreover, regulatory capacity of some tRFs has been observed; Zhou et al., for example, showed that a fragment excised from 5’ end of tRNA-Glu regulates BCAR3 expression in ovarian cancer(19). It is now clear that ncRNA-derived miRNA-like fragments are precisely processed out of various types of ncRNA transcripts, and that this processing is evolutionarily conserved across species(10–13).

While an increasing body of evidence suggests specifically excised sncRNA fragments from an array of ncRNAs exist and are functionally relevant, there are currently no Web-based, user-friendly resources that offer comprehensive sncRNA fragment profiling and discovery, functional prediction, and the identification of significant differential expressions of fragments among datasets. To address this gap we present SURFR. SURFR refers to our <u>S</u>hort <u>U</u>ncharacterized <u>R</u>NA_<u>F</u>ragment <u>R</u>ecognition tool that identifies all miRNA, snoRNA, and tRNA fragments (as well as fragments from all other ncRNAs annotated in Ensembl) specifically excised in a given transcriptome provided as either a raw user-generated RNA-Seq dataset or NCBI SRR file identifier. In addition, SURFR can also compare individual fragment expressions among as many as 30 distinct datasets (as well as compare the expressions of full length (non-fragmented) sncRNAs).

### SURFR Features

- Identifies fragments specifically excised from all miRNAs, tRNAs, rRNAs, scaRNAs, scRNAs, snoRNAs, sRNAs, vault RNAs, and any other ncRNAs annotated in the current Ensembl assembly(20) in individual small RNA-Seq datasets.
- Ten files can be processed at once then up to 30 individual files compared after processing for ncRNA fragment differential expression analysis.
- SURFR can also determine and compare the expressions of all full length (non-fragmented) sncRNAs in a given transcriptome.
- SURFR results are stored on the server indefinitely, protected by powerful state-of-the-art cryptographic algorithms, and can be instantly recalled by the user via entering their session key in the “Get Results” tab on the SURFR home page.
- OmniSearch-based miRNA analysis of annotated miRNAs(21).
- Direct, intuitive ncRNA visualization of individual ncRNA fragmentation.
- Easily downloadable Excel files of results from a single RNA-Seq file and/or comparisons among files. These files can be filtered (if desired) and list clearly defined, readily understandable, pertinent data (e.g., fragment expression, host gene links, and the exact fragment sequence excised).
- Contains prepopulated ncRNA databases allowing the identification of ncRNA fragments and/or ncRNA expressions in 440 unique animal, plant, fungal, protist, and bacterial species.

In addition, SURFR RNA fragment calls require considerably less processing time than previous ncRNA fragment identification pipelines for two principle reasons. We have: (1) developed a novel alignment strategy significantly faster than traditional methods (e.g., BLAST(22)) and (2) designed a novel method to locate the start and end positions of an ncRNA fragment using wavelets. Full details of these novel computational methodologies are described in length in **Supplemental Information File 2**.

### SURFR Workflow

### SURFR Input

Under “Use SURFR”, the user first selects the organism corresponding to the sequences. SURFR small RNA databases have been prepopulated for 440 species including 286 metazoans, 62 plants, and 92 other fungi, protists, and bacteria. As indicated in **Figure 1**, the user then provides one to ten small RNA sequencing datasets as input. These datasets can be all uploaded directly by the user, or all downloaded from the NCBI SRA database(23) by entering SRA IDs (e.g., SRR6495855, SRR4217122), or any combination thereof (for example, three datasets uploaded by the user along with seven datasets downloaded from the NCBI SRA database). Importantly, a major strength of SURFR is that users can upload most raw small RNA-Seq files directly as original, unmodified, compressed FASTQ files (as provided by commercial sequencers) with absolutely no preprocessing and with no specifics about library generation, linkers, or oligonucleotides required. Allowable formats for uploading are uncompressed, standard FASTA or FASTQ files or any major compression of either.

**Figure 1.**
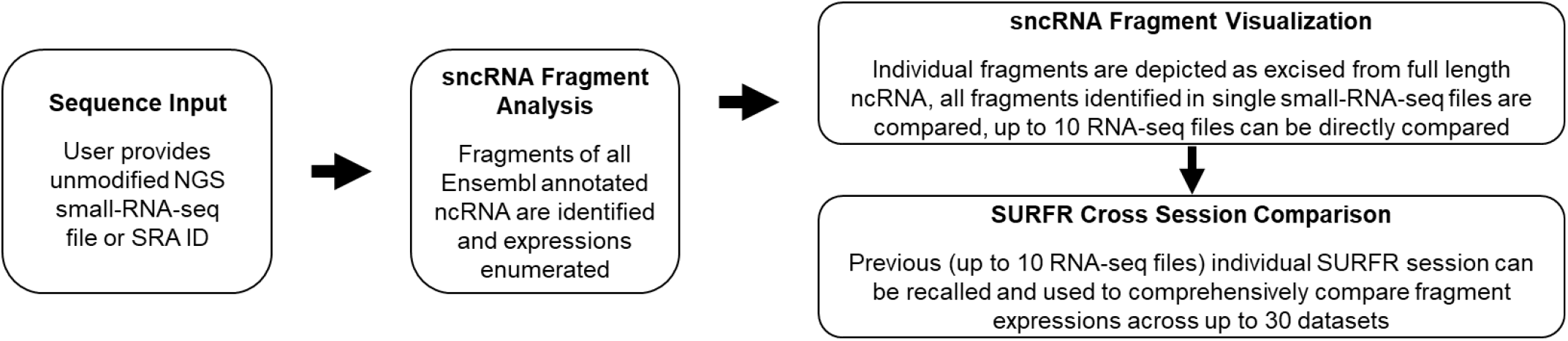
SURFR workflow. Sequence Input (left). The user provides up to ten unmodified small RNA-Seq datasets as input. These datasets can all be uploaded directly by the user or downloaded from the NCBI SRA database by entering SRA IDs. sncRNA Fragment Analysis (middle). SURFR identifies all ncRNA fragments (both annotated and novel) and their expressions in up to ten datasets per session. sncRNA Fragment Visualization (top right). Graphics of individual host ncRNAs and the fragments excised (along with the expressions at each nt position) are provided. In addition, tables comparing the expressions of all fragments within individual datasets and comparing fragment expressions across all datasets are generated. SURFR Cross Section Comparison (bottom right). The user can comprehensively compare all fragment expressions identified in up to 30 individual datasets by entering multiple SURFR session IDs from separate analyses.

### SURFR Output

After the user uploads/specifies the small RNA-Seq datasets and clicks the “Let’s SURF” button, the browser is automatically redirected to a report page, progress indicators for each uploaded dataset are provided under the “Click Here To Choose Your File” drop down menu at the top of the page (**Figure 2A**) with individual datasets having completed analysis indicated by a checkmark. Following completion of analysis, results for the individual file selected are then displayed on the report page and organized into several sections (**Figure 2**).

**Figure 2.**
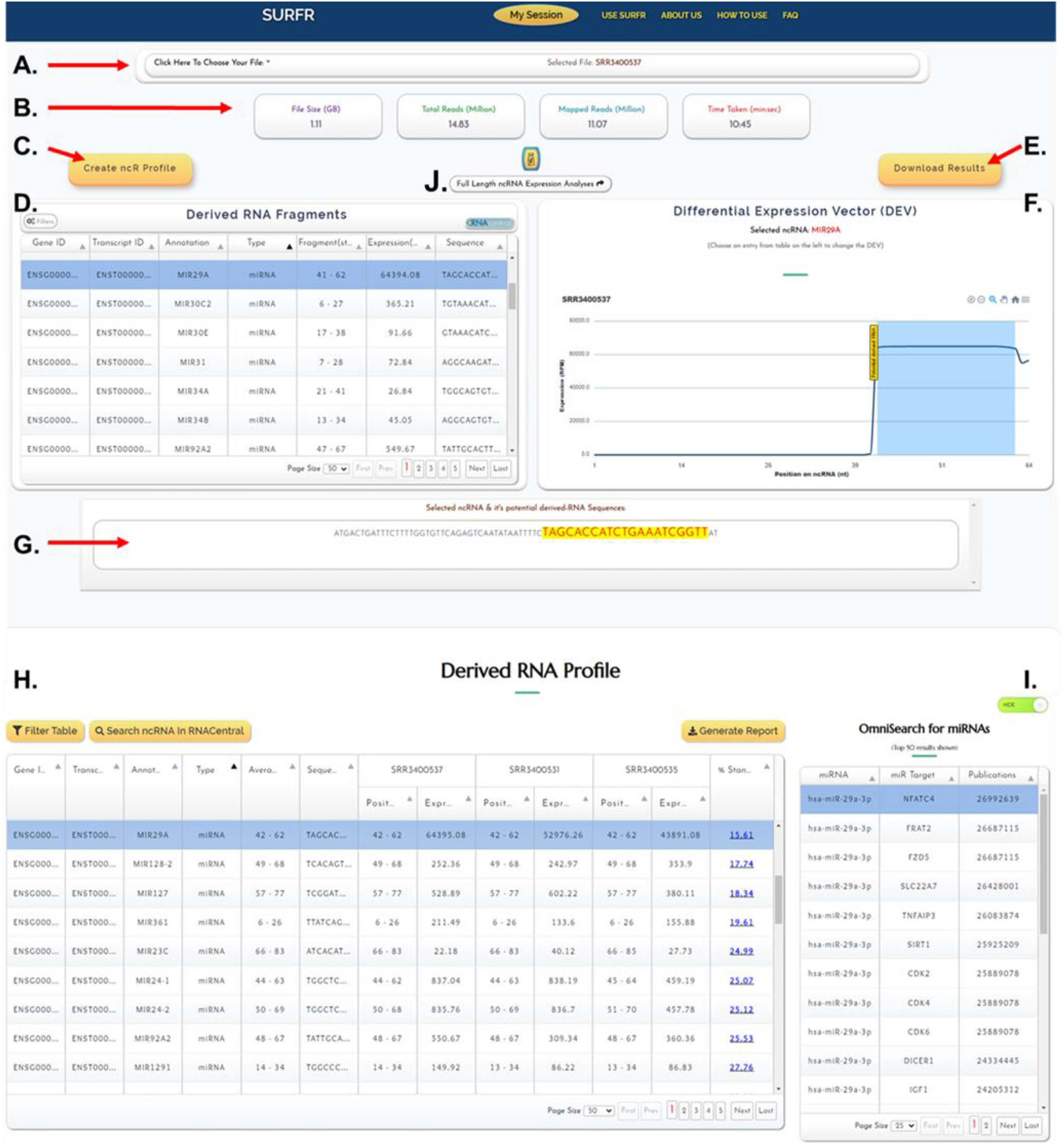
SURFR report page. SURFR report example. (**A**) The “Click Here To Choose Your File” drop-down menu for selecting individual RNA-Seq files. (**B**) A summary of the overall composition of the selected small RNA-Seq dataset. (**C**) The “Create ncR Profile” button automatically populates the derived RNA Profile section at the bottom of the page. (**D**) The “Derived RNA Fragments” window detailing each fragment identified in the individual, selected small RNA-Seq dataset. (**E**) The user can download an Excel file detailing the full set of information presented in the “Derived RNA Fragments” window by pressing the “Download Results” button. (**F**) The “Differential Expression Vector (DEV)” window illustrates each nucleotide within a host gene and indicates the fragment called with a blue rectangle. The x-axis represents the position in the ncRNA selected (e.g., miR-29a), and the y-axis depicts the expression levels of the ncRNA at each position. (**G**) The “Selected ncRNA & Called RNA Fragment Sequences” window illustrates the full length host ncRNA (miR-29a) highlighting the SURFR-called fragment in yellow. (**H**) The “Derived RNA Profile” window details each fragment identified in any of the analyzed small RNA-Seq datasets and compares fragment expressions across samples. (**I**) The “OmniSearch for miRNAs” window lists the top 50 OmniSearch entries (reported targets and PubMed publications) for an individual miRNA selected in the “Derived RNA Profile” window. (**J**) The “Full Length ncRNA Expression Analyses” button in the upper center of the results page redirects the user to a SURFR window detailing the expressions of all full length sncRNAs in the provided datasets.

A summary of the overall composition of the selected small RNA-Seq dataset, including the file size, total number of reads, number of mapped reads, and time taken for analysis is included just below the file selection window at the top of the page (**Figure 2B**).

The user can compare fragment expressions across all datasets by pressing the “Create ncR Profile” button that automatically populates the derived RNA Profile section at the bottom of the page (**Figure 2C**).

The “Derived RNA Fragments” window (**Figure 2D**) details the Ensembl Gene ID, Ensembl Transcript ID, gene annotation (name), the type of gene a fragment was excised from, the start and end positions of a fragment within its host gene, the expression of a fragment in reads per million (RPM), and the nucleotide sequence for each fragment identified in the individual, selected small RNA-Seq dataset. The “Derived RNA Fragments” window is an interactive table that allows users to view, sort, and filter small RNA fragments based on any column value. Users can also view host gene information available at the RNAcentral browser by selecting a fragment in the table and then clicking the “RNAcentral” button on the toolbar(24).

The user can download an Excel file detailing the full set of information presented in the “Derived RNA Fragments” window (**Figure 2D**) for each fragment identified in the individual, selected small RNA-Seq dataset by pressing the “Download Results” button (**Figure 2E**). An Excel file containing the derived RNA fragment information in its entirety will be automatically downloaded to the user’s computer (**Figure 3**).

**Figure 3.**
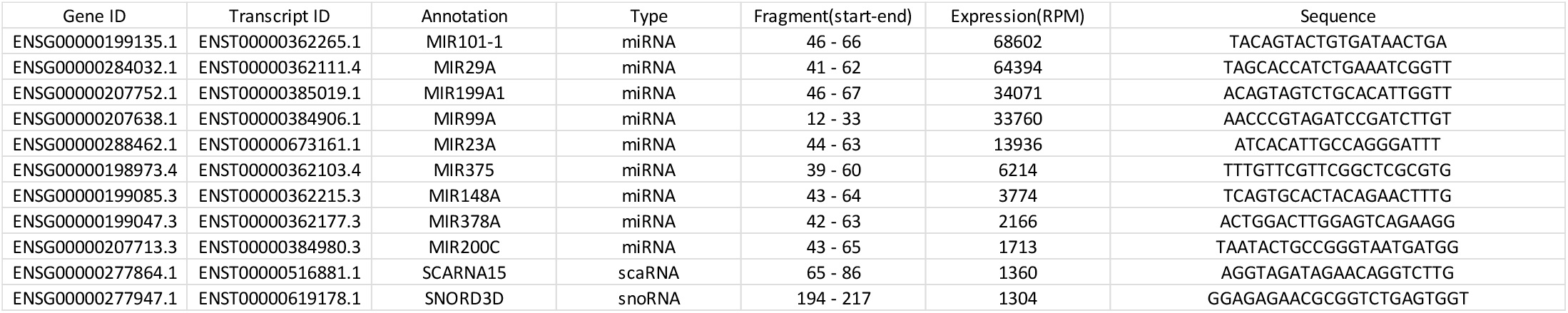
Derived RNA Fragments “Download Results” File. The first few rows of an example “Download Results” Excel file detailing the full set of information presented in the “Derived RNA Fragments” window: Ensembl “Gene ID”, Ensembl “Transcript ID”, gene “Annotation” (name), the “Type” of gene a fragment was excised from, the start and end positions of a fragment within its host gene, the expression of a fragment in reads per million (RPM), and the nucleotide “Sequence” for each fragment identified in the selected small RNA-Seq dataset.

The “Differential Expression Vector (DEV)” window (**Figure 2F**) details the expressions of each nucleotide within a host gene and indicates the fragment called with a blue rectangle. The x-axis in the graph shown in Figure 2F represents the position in the ncRNA selected (miR-29a), and the y-axis represents the expression levels of the ncRNA at each position. The user can also interactively view the expression at each individual nucleotide by panning over the image, zoom in or out using the buttons on the top right, and/or download DEV image files and an Excel file detailing expression at each nucleotide by selecting the menu button on the top right of the window.

The “Selected ncRNA & Called RNA Fragment Sequences” window (**Figure 2G**) illustrates the full length host ncRNA highlighting the SURFR-called fragment in yellow just as depicted in the preceding DEV window (**Figure 2F**).

The “Derived RNA Profile” window (**Figure 2H**) details the Ensembl Gene ID, Ensembl Transcript ID, gene annotation (name), the type of gene a fragment was excised from, the average start and end positions of a fragment within its host gene (To be considered the same fragment start and stop positions had to agree within 5 nts.) with corresponding nucleotide sequence for each “average” fragment listed, the start and end positions of a fragment within its host gene along with the fragment’s expression (RPM) in each individual small RNA-Seq dataset, and finally, the % standard deviation of the expression of individual fragments(20). Importantly, the full list of all fragments identified in any of the datasets is presented. The “Derived RNA Profile” window is an interactive table that allows users to view, sort, and filter small RNA fragments based on any column value. Users can also view host gene information available at the RNAcentral browser(24) by selecting a fragment in the table and then clicking the “Search ncRNA In RNAcentral” button on the toolbar.

The user can also download an Excel file detailing the full set of information presented in the “Derived RNA Profile” window by pressing the “Generate Report” button at the top right of the window. An Excel file containing the derived RNA profile information in its entirety will be automatically downloaded to the user’s computer (**Figure 4**). In addition, Excel file reports can be downloaded following the application of specific filters in the “Derived RNA Profile” window (e.g., only snoRNA fragments can be included or excluded).

**Figure 4.**
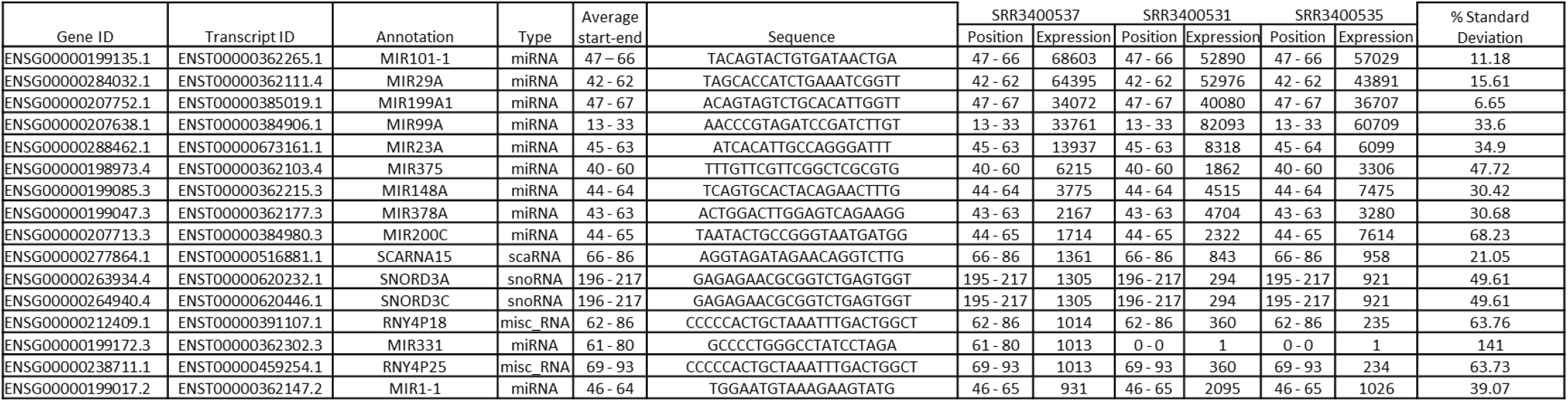
Derived RNA Profile “Generate Report” File. The first few rows of an example “**Generate Report**” Excel file detailing the full set of information presented in the “Derived RNA Profile” window.

The “OmniSearch for miRNAs” window (**Figure 2I**) returns the top 50 OmniSearch entries(21) (reported targets and PubMed entries) for an individual miRNA selected in the preceding “Derived RNA Profile” window.

And finally, when desired, the “Full Length ncRNA Expression Analyses” button (**Figure 2J**) redirects the user to a SURFR window detailing the expressions of all full length sncRNAs in the provided datasets regardless of fragmentation. Importantly, all pertinent features (e.g. expression table downloads) described above are similarly available for full length sncRNA analyses via this resource.

### SURFR Example Use/Case Study

SURFR allows users to profile and compare the expressions of sncRNA fragments (both annotated and novel) across multiple small RNA-Seq experiments in order to identify the top sncRNA fragments significantly differentially expressed in a particular disease, tissue, developmental stage, etc.

Our group’s interest in fragments excised from ncRNAs other than miRNAs initially arose from an attempt to identify novel miRNA contributors to breast cancer(12). For this work, we performed small RNA sequencing on several breast cancer cells lines, and while we failed to identify any (traditional) miRNAs of interest, we did identify a snoRNA fragment (we deemed sdRNA-93) that was specifically and significantly overexpressed in MDA-MB-231 cells-a widely studied model of a highly invasive and metastatic human cancer. Next, as we found sdRNA-93 to be significantly overexpressed in these cells (≥75x compared to controls), we decided to determine if sdRNA-93 functionally contributed to the malignant phenotype. Stringently testing sdRNA-93 inhibitors and mimics in MDA-MB-231 cells across multiple time points revealed that sdRNA-93 gain-and loss-of-function showed profound effects on invasion within standard matrix-based (matrigel) chemoattractant assays. Remarkably, sdRNA-93 loss-of-function reduced cell invasion by >90% at 48 hours compared to control cells, whereas sdRNA-93 gain-of-function enhanced cell invasion by >100%. Thus, we showed a single sdRNA (sdRNA-93) strongly selectively regulates invasion of MDA-MB-231s. These findings link a specific sdRNA (sdRNA-93) to an aggressive malignant phenotype (invasion) within an established cancer cell model that is widely used to study invasive behavior. We next employed a BLAST-based methodology to determine sdRNA-93 expressions across small RNA-Seq datasets corresponding to 115 unique breast cancer patients and detected strong overexpression of sdRNA-93 in 92.8% of tumors classified as Luminal B Her2+, compared to normal tissue controls (extremely low expression) and other breast cancer subtypes (modest expression levels of 30-40%). Thus, this work represented the first evidence demonstrating that sdRNAs that regulate specific malignant properties are differentially expressed within divergent molecular subtypes of human breast cancer(12).

Importantly, our initial BLAST-based identification of sdRNA-93 as being significantly overexpressed in MDA-MB-231 cells was highly labor intensive taking days to complete. In contrast, when we uploaded our original unmodified FASTQ sequencing files to SURFR, sdRNA93 was readily identified as the most highly differentially expressed snoRNA fragment between our two cancer cell lines taking just 7.9 minutes (**Figure 5**).

**Figure 5.**
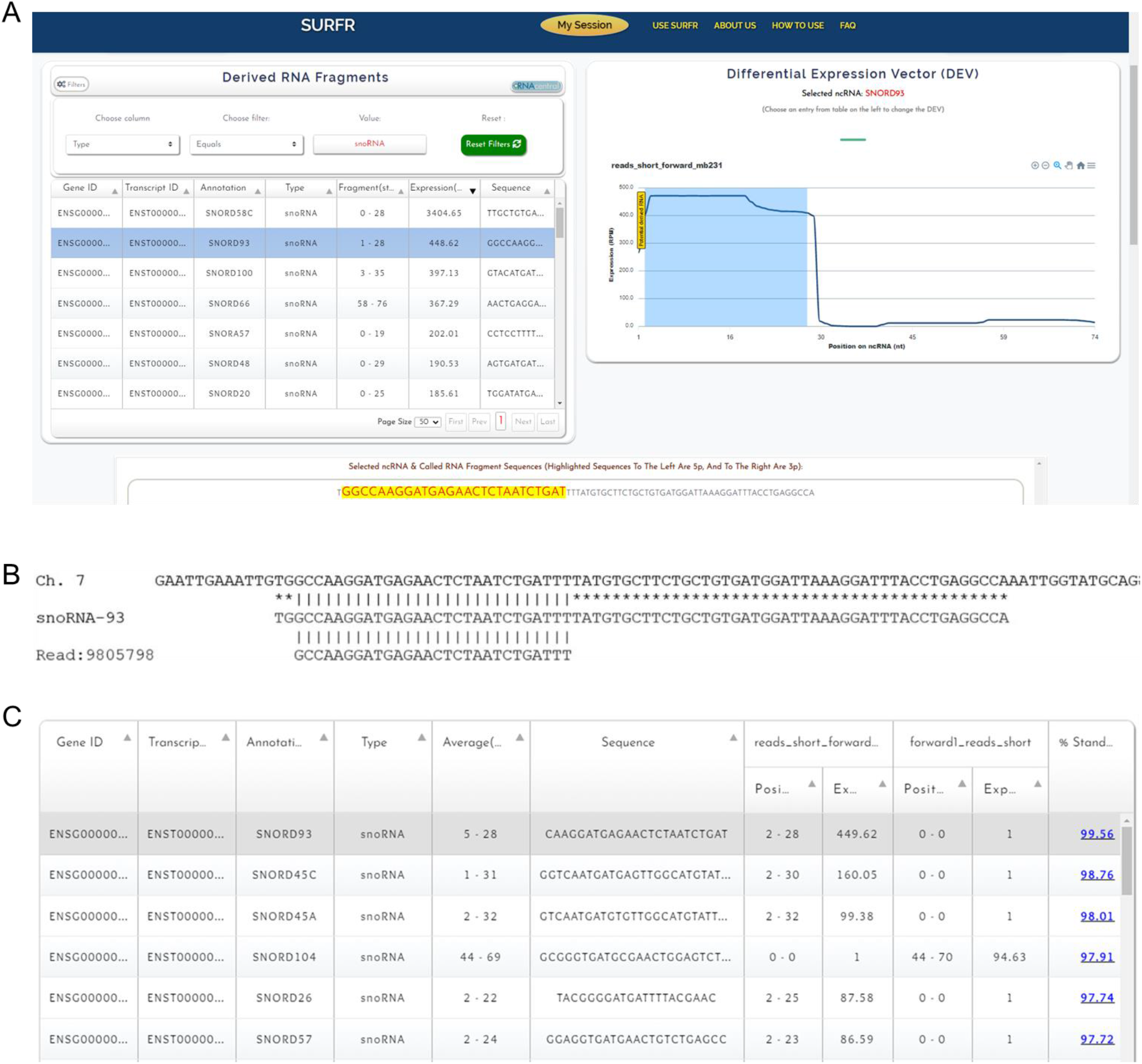
SURFR identification of sdRNA-93. (**A**) “Derived RNA Fragments” window showing SNORD93 derived sdRNA-93 was identified as the second most highly expressed sdRNA in the highly invasive breast cancer cell line MDA-MB-231. (**B**) Alignment among the human genome (GRCh38 Ch7:22856601:22856699:1) (top), snoRNA-93 (ENSG00000221740) (middle), and next generation small RNA sequence read (bottom) obtained by Illumina sequencing of MDA-MB-231 RNA as originally described in(12). All sequences are in the 5′ to 3′ direction. An asterisk indicates base identity between the snoRNA and genome. Vertical lines indicate identity across all three sequences. (**C**) “Derived RNA Profile” window comparing small RNA-Seq results for MCF-7 and MDA-MB-231 cells. Note SNORD93 derived sdRNA-93 was identified as the most significantly differentially expressed sdRNA between the weakly and highly invasive breast cancer cell lines.

### SURFR Comparison to other Existing Tools

Numerous characterizations of significant regulatory roles for sncRNA fragments excised from various types of ncRNAs other than miRNAs have now been reported(10–13). As new high-throughput small RNA sequencing strategies(25) continue to make small RNA-Seq faster and less expensive, there is a clear need for tools capable of digesting large amounts of small RNA-Seq data in order to detect and characterize all small RNA genes including specifically-excised small RNA fragments. Most existing tools (e.g., miRDeep(26), miRSpring(27), miRanalyzer(28), etc.) focus almost exclusively on miRNAs and/or only evaluate existing sncRNA annotations and are not capable of fully defining small RNA-Seq ncRNA fragment profiles and differences among these datasets (sRNAnalyzer(29), Oasis2.0(30), SPAR(31), etc.). That said, most existing tools capable of characterizing novel ncRNA fragments and their expressions, such as FlaiMapper(32), SPORTS(33), and DEUS(34), require fairly extensive computational expertise for utilization, support only pre-aligned file inputs (BAM), and/or require standalone installation (**Table 1**). As such, we have designed SURFR to address the need for a user-friendly, Web-based, comprehensive small RNA fragment tool requiring no computational expertise to utilize. In stark contrast to most existing platforms, SURFR identifies fragments excised from all types of ncRNAs annotated in Ensembl(20) in a given transcriptome provided as either a raw user-generated RNA-Seq dataset or NCBI SRA file. In addition, SURFR can compare individual fragment expressions among as many as 30 distinct datasets, and we have included ncRNA databases for 440 unique animal, plant, fungal, protist, and bacterial species. Importantly, there are currently no Web-based, user-friendly resources that offer comprehensive sncRNA fragment profiling and discovery, functional prediction, and the identification of significant differential expressions among datasets comparable to SURFR. Although two platforms, sRNA toolbox(35) and sRNAtools(36), do offer many of SURFR’s features, SURFR distinguishes itself by providing significantly more intuitive, versatile, and user friendly results generated in less than 10% of the time required for data upload and processing by these tools. That said, because SURFR was developed specifically for ncRNA fragment identification, it does not provide expression analysis for full length ncRNAs.

**Table 1.** SncRNA analysis platform feature comparison. Various features offered by SURFR were compared to other existing tools including sRNA toolbox(35), Oasis2.0(30), sRNAtools(36), CPSS2.0(37), SPAR(31), sRNAnalyzer, SPORTS1.0(33), DEUS(34), FlaiMapper(32), and featureCounts(38). Features examined were: “Online,” if tool is available online; “Input,” form of input RNA-Seq dataset-either raw (direct NGS output) or pre-processed (e.g., requires BAM file); “Clear, User-friendly Results/Output,” if interactive and user-friendly results are generated directly; “Library Oligo Sequences Req,” if user knowledge of NGS oligo sequences is required; “TCGA, SRA, GEO, or Encode Input,” if publically available RNA-Seq datasets can be specified for examination based on identifier alone; “Known Full Length sncRNA Expressions,” detection and quantification of known sncRNAs; “Novel Full Length sncRNA Expressions,” detection and quantification of novel sncRNAs; “Novel sncRNA Fragment Discovery,” detection and quantification of novel ncRNA fragments; “Differential Expression,” ability of the tool to integrate expression data from multiple files (“sRNAde” denotes that expression analyses can be performed in parallel); and “Species,” number of species available for analysis. “user” denotes that the tool has the capacity to perform given task however requires additional user input or user-directed change to program’s code and/or advanced settings.

| PROGRAM | Online | Input | Clear, User-friendly Results / Output | Library Oligo Sequences Req? | TCGA, SRA, GEO, or Encode Input | Known Full Length sncRNA Expressions | Novel Full Length sncRNA Discovery | Novel sncRNA Fragment Discovery | Differential Expression | Species |
| --- | --- | --- | --- | --- | --- | --- | --- | --- | --- | --- |
| SURFR | yes | raw | yes | no | yes | no | no | yes | yes | 440 |
| sRNA toolbox | yes | raw | no | yes | yes | yes | yes | no | sRNAde | 175 |
| Oasis2.0 | yes | raw | yes | yes | no | yes | yes | no | yes | 14 |
| sRNAtools | yes | raw | no | yes | yes | yes | yes | yes | yes | 21 |
| CPSS2.0 | yes | pre-processed | yes | yes | no | yes | yes | no | yes | 48 |
| SPAR | yes | pre-processed | yes | yes | yes | yes | yes | no | yes | 2 |
| sRNAAnalyzer | no | raw | no | yes | no | yes | no | no | no | user |
| SPORTS1.0 | no | raw | no | yes | yes | yes | user | tRF, rRF only | yes | 68 |
| DEUS | no | raw | no | yes | no | yes | yes | user | yes | user |
| FlaiMapper | no | pre-processed | no | yes | no | no | no | yes | no | user |
| featureCounts | no | pre-processed | no | yes | no | user | no | user | no | user |

Notably, as a verification of SURFR’s accuracy, we recreated an analysis of ten prostate cancer small RNA-Seq files previously performed using FlaiMapper(39). Importantly, FlaiMapper-based ncRNA fragment discovery of these ten files originally identified 147 snoRNA-derived fragments that were 18 to 35 nt in length and expressed at > 10 RPM. Similarly, SURFR analysis of the same files identified 110 snoRNA-derived fragments expressed at > 10 RPM, and strikingly, 104 of these fragments were nearly identically identified (+/-2 nts) by both methods. Notably, we find the majority of the FlaiMapper-identified sdRNA fragments not present in the SURFR calls were excluded based on SURFR’s 100% sequence identity requirement (in contrast to FlaiMapper’s 2 nt mismatch allowance).

### SECTION 2. SALTS Tool for Long non-coding RNA Analysis: LAGOOn

ncRNAs longer than 200 nt in length are known as long ncRNAs (lncRNAs). This distinction, while somewhat arbitrary and based on technical aspects of RNA isolation methods, serves to distinguish lncRNAs from miRNAs and other sncRNAs. lncRNA loci are present in large numbers in eukaryotic genomes typically comparable to or exceeding that of protein coding genes. Many lncRNAs possess features reminiscent of protein-coding genes, such as having a 5′ cap and undergoing alternative splicing(40). In fact, many lncRNA genes have two or more exons(40), and about 60% of lncRNAs have polyA+ tails. In addition, although numerous long intergenic RNAs (lincRNAs)(41) including eRNAs from gene-distal enhancers have recently been reported(42), the majority of lncRNA genes identified to date are located within 10 kb of protein-coding genes and typically found to be antisense to coding genes or intronic(43). That said, many lncRNAs are expressed at relatively low levels in highly specific cell types(40) both explaining why the majority of lncRNAs were thought to be “transcriptional noise” until quite recently and also representing perhaps the single largest challenge in terms of lncRNA discovery and characterization.

NGS has now identified tens of thousands of lncRNA loci in humans alone with the number of lncRNAs linked to human diseases quickly increasing. That said, lncRNA functionality is highly contentious, and the number of experimentally characterized and / or disease-associated lncRNAs remain in the low hundreds, or ≤1% of identified loci(44). This has led to a burgeoning focus on elucidating the molecular mechanisms that underlie lncRNA functions(45). Although only a minority of identified lncRNAs have been functionally characterized, several distinct modes of action for lncRNAs have now been described, including functioning as signals, decoys, scaffolds, guides, enhancer RNAs, and short peptide messages(46)(47). Importantly, however, there are currently no Web-based, user-friendly resources that offer comprehensive lncRNA profiling, functional prediction, and the identification of significant differential expressions among datasets. To address this gap we present LAGOOn. LAGOOn refers to our <u>L</u>ong-noncoding and <u>A</u>ntisense <u>G</u>ene <u>O</u>ccurrence and <u>On</u>tology tool that identifies all lncRNAs expressed in a given human transcriptome from either a user-provided RNA-Seq dataset or publically available SRA file(23). In addition, LAGOOn can also compare lncRNA expressions among datasets and predict likely functional roles for individual lncRNAs.

### LAGOOn Features

- Direct, intuitive visualization of significant lncRNA expressions. Determines the expressions of all lncRNAs annotated in the current Ensembl assembly(20) in individual human RNA-Seq datasets.
- Identifies differentially utilized lncRNA exons.
- Up to three files can be processed at once then up to 15 individual files compared after processing for lncRNA differential expression analysis.
- LAGOOn results are stored on the server indefinitely, protected by powerful state-of-the-art cryptographic algorithms, and can be instantly recalled by entering a previous session key in “Access Your Results” on the LAGOOn home page.
- Easily downloadable Excel files of results profiling a single RNA-Seq file and/or comparisons among various files. These files can be filtered (if desired) and list clearly defined, readily understandable, and pertinent data (e.g., expression, lncRNA Ensembl ID, etc.).
- Detailed, comprehensive lncRNA functional prediction detailing:

- If a lncRNA serves as a host for a sncRNA(45).
- Significant potentials for a lncRNA to serve as a specific miRNA sponge(48).
- All overlaps between a given lncRNA and annotated enhancers(49).
- Significant potentials for lncRNAs to serve as naturally occurring antisense silencers for genes located on the strand opposite to themselves(50).
- Associations between individual lncRNAs and ribosomes suggesting microprotein production(51).

Importantly, LAGOOn is the first Web-based, user-friendly resource that offers real-time lncRNA profiling, the identification of significant differential expressions among datasets, and an array of functional prediction assessments beyond standard mRNA interaction characterizations. Full details of these novel computational methodologies are described in length in **Supplemental Information File 3**.

### LAGOOn Workflow

### LAGOOn Input

As summarized in **Figure 6**, after selecting “Start New Analysis” on the LAGOOn homepage, the browser is redirected to the “Data Transfer Options” page where the user provides one or two RNA sequencing datasets as input and is given the chance to provide an optional, additional input, i.e., a Ribo-Seq dataset for determining microprotein coding potentials. These datasets can all be uploaded directly by the user, or all downloaded from the NCBI SRA database(23) by entering SRA IDs (e.g., SRR9729388, SRR6290085), or any combination thereof. Importantly, a major strength of LAGOOn is that users can upload most raw RNA-Seq files directly as original, unmodified, compressed FASTQ files (as provided by commercial sequencers) with absolutely no preprocessing and with no specifics about library generation, linkers, or oligonucleotides required. There is no limit on the size of SRA files whereas individual user uploaded files are limited to 18 GB regardless of format meaning extremely large sequencing files exceeding even this size can be converted to FASTA format then compressed prior to being uploaded if necessary. Allowable uploaded formats are uncompressed, standard FASTA or FASTQ files or any major compression of either. In addition to this, the LAGOOn homepage provides links to: (1) “Access Your Results” where users can retrieve results from previous sessions via providing a session key and then compare results from up to five separate sessions. (2) “LAGOOn Search” where users can obtain detailed, comprehensive functional predictions for individual lncRNAs. And, (3) “Download Our Databases” where users can download databases containing all the lncRNAs and/or lncRNA exons employed by LAGOOn.

**Figure 6.**
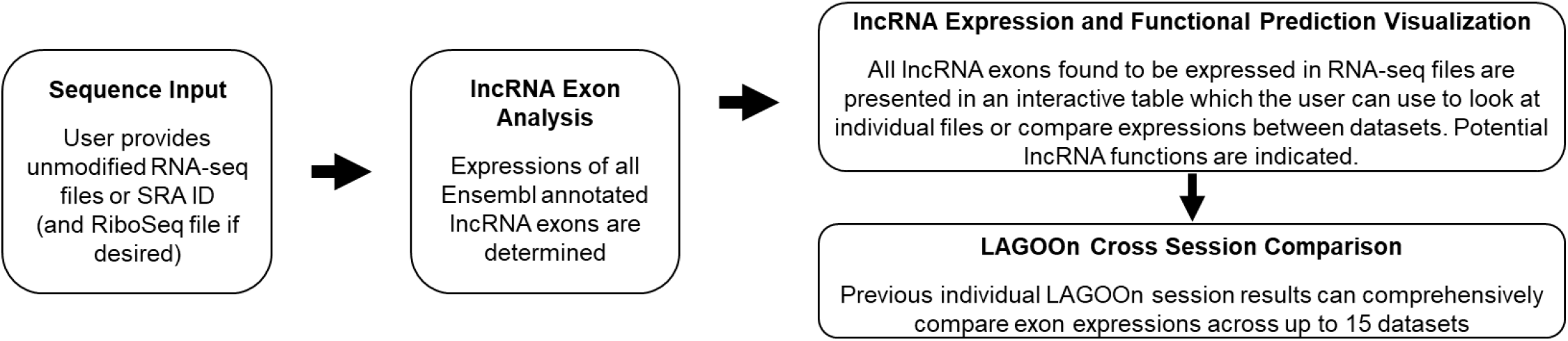
LAGOOn workflow. Sequence Input (left). The user provides up to two unmodified RNA-Seq files and one Ribo-Seq dataset (optional) as input. These datasets can all be uploaded directly by the user or downloaded from the NCBI SRA database by entering SRA IDs. lncRNA Exon Analysis (middle). LAGOOn enumerates all annotated lncRNA expressions in up to three datasets per session. lncRNA Expression and Functional Prediction Visualization (top right). An interactive table is generated comparing the expressions of all exons within individual datasets and comparing exon expressions across all datasets. Tables indicating putative lncRNA functions are also depicted. LAGOOn Cross Section Comparison (bottom right). The user can comprehensively compare all exon expressions identified in up to 15 individual datasets by entering multiple LAGOOn session IDs from separate analyses.

### LAGOOn Output

After the user uploads/specifies the RNA-Seq datasets, the browser is automatically redirected to the LAGOOn report page (**Figure 7**). Initially, a summary of the size and composition of individual RNA-Seq datasets, the number of lncRNAs expressed in a dataset, and the top ten most highly expressed lncRNAs in the specified dataset are shown. Following selection of either one or all of the RNA-Seq files and the Ribo-Seq file (if included) analyzed from the file selection toolbar (**Figure 7A**), results for the file(s) selected are then displayed on the report page under the “Results” tab (**Figure 7B**), and organized into several distinct sections.

**Figure 7.**
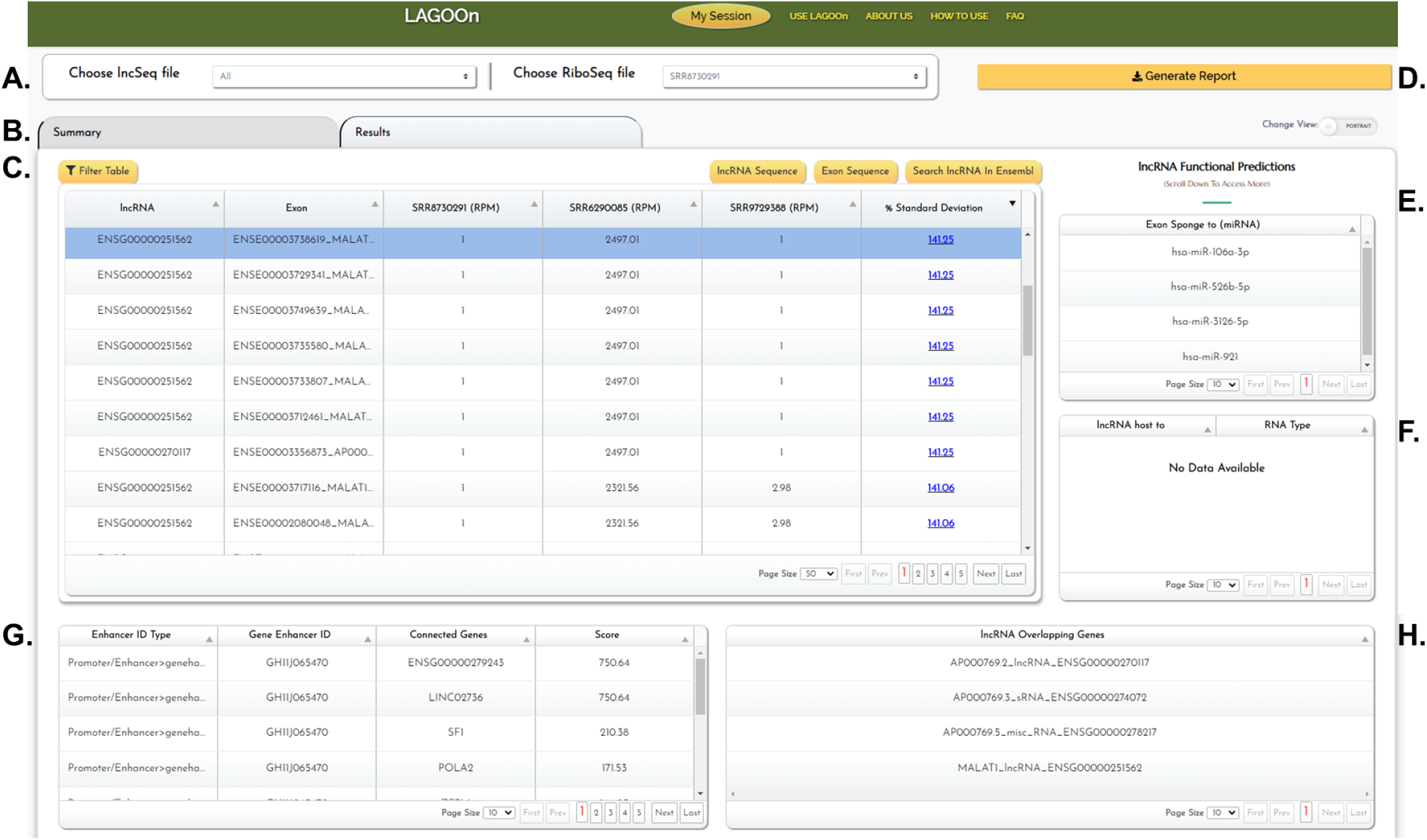
LAGOOn report page. LAGOOn report example. (**A**) The file selection toolbar contains drop-down menus for selecting individual RNA-Seq and Ribo-Seq files. (**B**) The toolbar allowing selection of either the “Summary” or “Results” tab. (**C**) The lncRNA expression window displays a filterable table of all lncRNA exons expressed in any of the user-provided files. Full length lncRNA sequence, individual exon sequence, or Ensembl lncRNA gene information is obtained by selecting an exon in the table and then clicking the “lncRNA Sequence,”“Exon Sequence,” or “Search lncRNA in Ensembl” button on the toolbar. (**D**) The “Generate Report” button creates and automatically downloads an Excel file detailing the full set of information presented in the expression table window. (**E**) The “Exon Sponge to (miRNA)” window lists all miRNA complementarities of ten base pairs or greater occurring within the selected lncRNA exon (**F**) The “lncRNA host to” window lists all full length ncRNAs contained in any of the selected lncRNA’s exons. (**G**) The “Enhancer” window lists all overlaps between a selected lncRNA and GeneHancer annotated enhancer (as well as genes with expression linked to individual enhancers). (**H**) The “lncRNA Overlapping Genes” window lists all genes even partially overlapping a lncRNA locus on either strand.

The table presented in **Figure 7C** details the Ensembl Gene ID, Ensembl Exon ID along with gene annotation (name), and expressions (RPM) of all lncRNA exons in each individual RNA-Seq dataset, and finally, the % standard deviation of the expression of individual exons(20). Importantly, the full list of all exons found to be expressed in any of the datasets is presented. In addition, the expression table is interactive and allows user to view, sort, and filter based on any column value by clicking the “Filter Table” button on the toolbar. Users can also obtain a full length lncRNA sequence, a specific exon sequence, or view the lncRNA gene information available at Ensembl by selecting an exon in the table and then clicking the “lncRNA Sequence,”“Exon Sequence,” or “Search lncRNA in Ensembl” button on the toolbar.

The user can also download an Excel file detailing the full set of information presented in the expression table window by pressing the “Generate Report” button at the top right of the window (**Figure 7D**). An Excel file containing the expression table window information in its entirety will be automatically downloaded to the user’s computer (**Figure 8**). In addition, refined Excel file reports can be downloaded following the application of specific filters (e.g., lncRNAs with RPM > 1 in the Ribo-Seq dataset).

**Figure 8.**
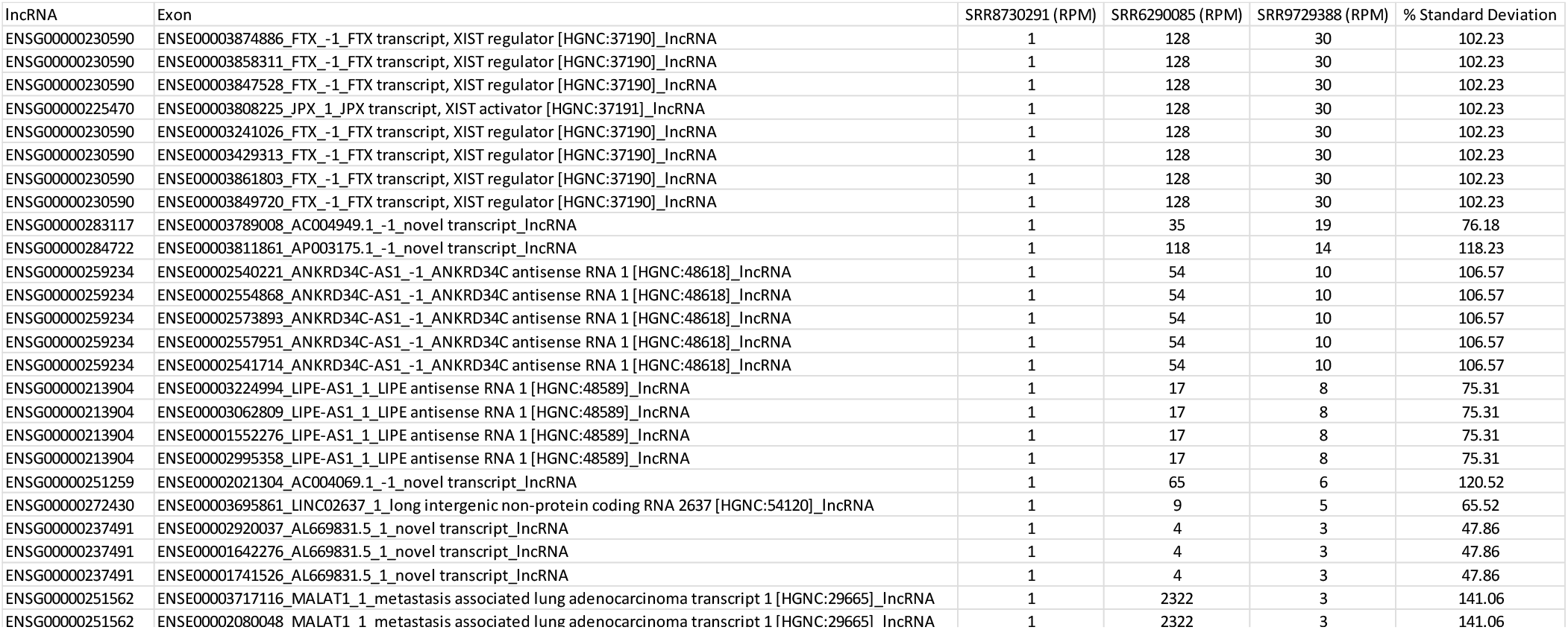
lncRNA expression table “Generate Report” File. The first few rows of an example “**Generate Report**” Excel file detailing the full set of information presented in the lncRNA expression window.

Finally, putative functional roles for lncRNAs/lncRNA exons selected in the expression table are depicted in **Figure 7E-H**. As lncRNAs frequently function as miRNA sponges that directly basepair with and effectively inactivate mature miRNAs(48), the “Exon Sponge to (miRNA)” window lists all miRNA complementarities of ten base pairs or greater occurring within the selected lncRNA exon (**Figure 7E**). Next, as numerous lncRNAs have been shown to encode sncRNAs (e.g., miRNAs and snoRNAs) in their exonic sequences, and sncRNA expression often relies on excision from the host lncRNA transcript(45), the “lncRNA host to” window lists all full length ncRNAs contained in any of the selected lncRNA’s exons (**Figure 7F**). In addition, as several lncRNAs have been reported to function through regulating the accessibility of transcriptional enhancers overlapping their genomic loci(49), all overlaps between a selected lncRNA and GeneHancer(52) annotated enhancer (and genes with expression linked to individual enhancers) are detailed in the “Enhancer” window (**Figure 7G**). And finally, in addition to lncRNA exonic sequences serving as sncRNA hosts, many sncRNAs are processed from lncRNA introns(45). Furthermore, many lncRNAs serve as naturally occurring antisense silencers of genes located on the strand opposite to themselves(50). For both of these reasons, as well as other potential regulatory relationships, all genes overlapping a lncRNA locus on either the positive or negative strand are detailed in the “lncRNA Overlapping Genes” window (**Figure 7H**). Importantly, a comprehensive report detailing each of the functional predictions is also available for individual lncRNAs by selecting the “LAGOOn Search” button on the homepage after entering a lncRNA Ensembl gene identifier. Notably, this search functionality does not require full LAGOOn analysis.

### LAGOOn Example Use/Case Study

ncRNAs are becoming major players in disease pathogenesis such as cancer. Metastasis Associated Lung Adenocarcinoma Transcript 1 (MALAT1) is a nuclear enriched lncRNA that is generally overexpressed in patient tumors and metastases. Overexpression of MALAT1 has been shown to be positively correlated with tumor progression and metastasis in a large number of tumor types including breast tumors. Furthermore, an earlier study evaluating breast cancer patient samples showed that MALAT1 expression is higher in breast tumors as compared to adjacent normal tissues (reviewed in (53)). As such we elected to compare lncRNA expressions in a breast cancer cell line (MDA-MB-231) RNA-Seq dataset (SRR12101868) with those of a human bone tissue RNA-Seq dataset (SRR12101882) in order to identify significantly differentially expressed lncRNAs and their putative functions, including screening a Ribo-Seq of the BRX-142 cell line (SRR12101882) established from circulating tumor cells collected from a woman with advanced HER2-negative breast cancer(54) for potential MALAT1 microprotein production.

Strikingly, the total time for download and analysis of these three NGS datasets by LAGOOn was only 3 min 52 sec. More importantly, however, LAGOOn identified MALAT1 as the most highly expressed lncRNA in MDA-MB-231 breast cancer cells (**Figure 9**). In agreement with previous demonstrations that MALAT-1 functions (in part) as a miR-145-5p sponge in numerous malignancies including breast cancer(55), LAGOOn identified MALAT1 as a probable miR-145-5p sponge (**Figure 9A**, top right). In addition, LAGOOn also found MALAT1 overlaps with, and may therefore potentially be involved in regulating, several distinct genomic enhancers and sncRNAs (**Figure 9A**, lower windows). Finally, similarly in agreement with previous analyses(56), LAGOOn also identified MALAT1 as one of three lncRNAs significantly represented in the BRX-142 cell Ribo-Seq dataset strongly suggesting MALAT1 encodes at least one micropeptide.

**Figure 9.**
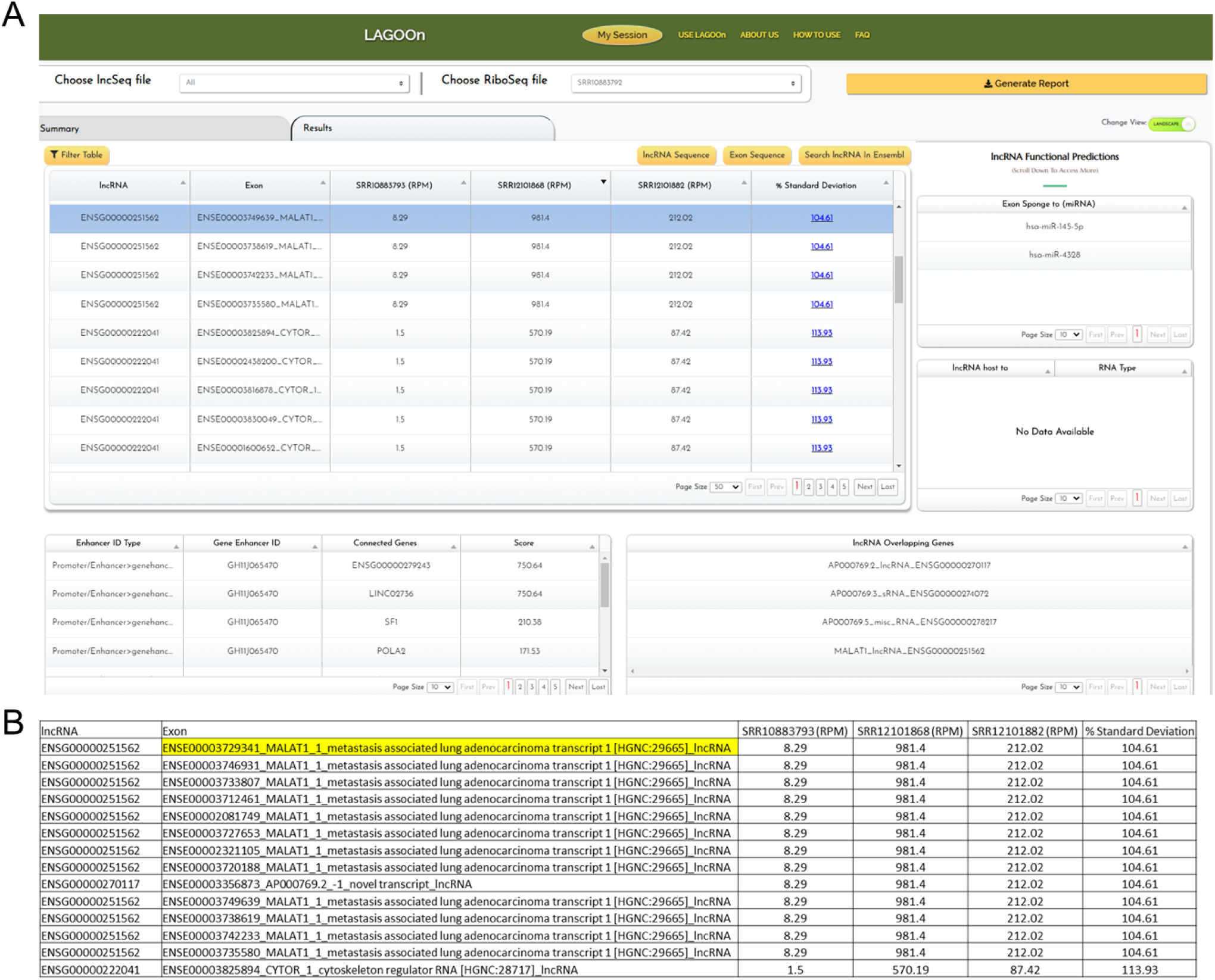
LAGOOn identification of MALAT1 overexpression in breast cancer. (**A**) The “Results” window showing MALAT1 was identified as the most highly expressed lncRNA in the highly invasive breast cancer cell line MDA-MB-231 (SRR12101868). (**B**) The “Generate Report” Excel file showing MALAT1 (yellow) was identified as the most highly expressed lncRNA in MDA-MB-231 cells. Both windows indicate MALAT1 is present in the breast cancer Ribo-Seq dataset (SRR10883792).

### LAGOOn Comparison to other Existing Tools

LncRNAs represent the largest single class of ncRNAs. However, unlike sncRNAs, which are thought to mostly function in gene regulation through complementary basepairing other RNAs, the mechanisms through which lncRNAs function are highly diverse. lncRNA relatively low expressions and tissue specificity have significantly hindered lncRNA discovery, our understanding of lncRNA regulations, and characterizations of lncRNA functional mechanisms to date(44)(45)(46)(47). That said, initiatives such as ENCODE(57), FANTOM(58), and GENCODE(40) have now predicted over 60,000 human lncRNAs and identified associations between many of these and specific diseases. Thus far, however, only a handful of these lncRNAs have been examined in the literature, with even fewer being assigned any specific mechanistic function.

Expression data often constitutes the first level of information of use in studying lncRNAs as differential expression analysis is clearly of value in prioritizing candidates for further examination. Differential expression, however, provides little in the way of functional insights. That said, the majority of computational platforms currently available are primarily aimed at either detecting and quantifying lncRNAs (e.g., lncRNA-screen(59), RNA-CODE(60), lncRScan(61), etc.) or predicting lncRNA:mRNA and/or lncRNA:protein interactions (e.g., PLAIDOH(62), LncRNA2Function(63), circlncRNAnet(64), etc.) (**Table 2**). In contrast, LAGOOn was designed to comprehensively evaluate lncRNA expression as well as the potential for lncRNAs to function through other characterized mechanisms including serving as sncRNA hosts, miRNA sponges, antisense RNAs, microprotein transcripts, and/or regulators of genomic enhancers (as well as providing links to predicted lncRNA:mRNA and/or lncRNA:protein interactions). In short, LAGOOn wholly distinguishes itself from available tools by filling a major gap in available lncRNA functional prediction platforms and eliminating the need of the user to switch platforms during the analysis process.

**Table 2.**
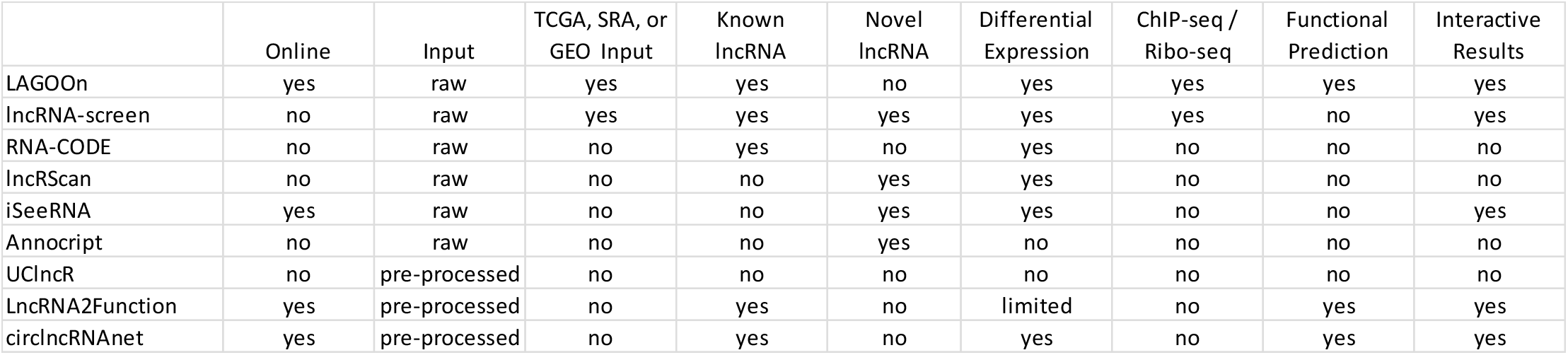
lncRNA analysis platform feature comparison. Various features offered by LAGOOn were compared to other existing tools including lncRNA-screen(59), RNA-CODE(60), lncRScan(61), iSeeRNA(65), Annocript(66), UClncR(67), LncRNA2Function(63), and circlncRNAnet(64). Features examined were: “Online”, if tool is available online; “Input”, form of input RNA-Seq dataset-either raw (direct NGS output) or pre-processed (e.g., requires BAM file); “TCGA, SRA, or GEO”, if publically available RNA-Seq datasets can be specified for examination based on identifier alone; “Known lncRNA”, detection and quantification of known lncRNAs; “Novel lncRNA”, detection and quantification of novel lncRNAs; “Differential Expression”, ability of the tool to integrate expression data from multiple files; “ChIP-Seq / Ribo-Seq”, if identified lncRNA occurrences in ChIP-Seq and/or Ribo-Seq datasets can be determined; “Functional Prediction”, if potential functional roles of identified lncRNAs are assessed; and “Interactive Results”, if interactive and user-friendly results are generated directly.

## DISCUSSION

Despite a mounting body of evidence supporting the physiological relevance of ncRNAs, most studies performed to date have focused primarily on proteins themselves or deciphering the pathways associated with annotated ncRNAs. Moreover, due to the perceived insurmountability of the sheer amount of data generated by NGS/TGS analyses, the full extent of regulatory networks created by ncRNAs often gets overlooked(68). In addition, whereas the cost of RNA-seq is now reasonable for most active research programs, tools necessary for the interpretation of these sequencing datasets typically require significant computational expertise and resources markedly hindering widespread utilization of these tools. As such, the necessity for development of real-time, user-friendly platforms capable of making the identification and characterization of the ncRNAome accessible to biologists lacking significant computational expertise becomes clear. In light of this, we have developed SALTS a highly accurate, super efficient, and extremely user-friendly one-stop shop for ncRNA transcriptomics. Notably, SALTS is accessed through an intuitive Web-based interface, can analyze either user-generated, standard NGS file uploads (e.g., FASTQ) or existing NCBI SRA datasets, and requires absolutely no dataset pre-processing or knowledge of library adapters/oligonucleotides. In short, SALTS constitutes the first publically available, Web-based, comprehensive ncRNA transcriptomic NGS analysis platform designed specifically for users with no computational background, providing a much needed, powerful new resource enabling more widespread ncRNA transcriptomic analysis.

That said, an array of platforms and pipelines, each geared towards a specific type of transcript/ncRNA class, have previously been developed. Regardless of the platform, the core of ncRNA transcriptome expression analysis consists of two main steps: transcript detection and expression quantification(1)(3). The first step in this process involves aligning, or mapping, the NGS reads to a reference sequence(s), which can be either ncRNA sequence library or an entire reference genome. Most standard pipelines use alignment programs such as Bowtie2(69), BWA(70), NCBI’s BLAST(22) or other implementations of existing alignment algorithms like Smith-Waterman (SW)(71), Needleman-Wunsch (NW)(72), and Burrows Wheeler Transform (BWT)(73). These aligners often differ in how alignment mis-matches and gaps are scored and as such need to be taken into account when dealing with data containing high sequence variability between the individual transcripts originating from the same genomic locus or between the reads and the reference. In the second step, aligned reads are further analyzed to determine the expression, or the number of reads assigned to individual loci or library entries. This step often includes or is followed by various statistical analysis to determine differential expression and/or variance between replicates (i.e., baySeq(74) or DESeq2(75)). That said, the strikingly high accuracy and efficiency achieved by our tools as compared to existing platforms is primarily due to a novel computational approach to RNA-Seq alignment and an innovative analysis based on Hilbert and Vector spaces developed in the course of this work. Brief overviews of the primary constructs critical to toolkit implementation are described below with more in-depth descriptions detailed in **Supplemental Information Files 2**and **3**.

### SALTS toolkit implementation

Of note, both SURFR and LAGOOn were developed into real-time processing systems using the following technology stack:

**Programming languages used**: Python 3.7, Visual C++ 2015, Erlang, JavaScript, PHP, and SPARQL.
**Database engines**: Mongo DB 4.4
**Servers**: Apache Web Server, 30+ background servers composed using Master-Worker model to parallelize the workload, and Apache Jena Fuseki.
**Other tools and supporting technologies**: Rabbit MQ, Flask, Redis, Vue JS, Dropzone JS, Apexcharts JS, Bootstrap 4, IBM Aspera, Axios JS, Moment JS, Tabulator, Matplotlib, NumPy, SciPy, and HTML5.
**Architecture**: Microservices.
**Hardware Specs**: Intel^®^ Xeon^®^ CPU-E5-2609 v4 @ 1.70GHz, 64GB RAM, 4TB Hard disk, Windows Server 2016.

### SURFR implementation

With SURFR, users with no computational background can quickly and easily analyze, visualize, and compare small RNA-Seq datasets in order to generate clear, informative results. With an interactive, user-friendly interface, SURFR is the first Web-based resource that provides users the ability to upload unmodified NGS datasets and/or provide SRA identifiers to perform comprehensive novel ncRNA and ncRNA fragment identifications and expression analyses in real-time. This is achieved through employing the following three key components: (1) <u>Hilbert Space (HS)</u>. In mathematics, a HS is an abstract vector space (with up to infinite dimensions) representing the current physical state of a continuous system routinely applied in Quantum mechanics. HSs are highly useful in describing the relationship among Vector spaces, Wavelets, and wave functions(76)(77). For our analyses, the term “Gene Expression” is considered a higher dimensional function representing the activity of the RNA across its length where, within a RNA, expression is represented using four vectors (for A, C, T, and G) and understood using HSs. (2) <u>MoVaK alignment.</u> Based on utilization of the aforementioned HSs, we introduced two new data structures, namely, Similarity Vectors (SVs) and Differential Expression Vectors (DEVs). MoVaK alignment combines SVs and DEVs to profile the exact transcriptomic activity of a given RNA-Seq dataset and then retrieves a HS for each RNA that is expressed in a sample. And (3) <u>SURFR algorithm.</u> By defining the changes in the gene expression using the above HS interpretation, we assign a wavelet function with scales of 18 to 38 to each sncRNA micro-like behavior, i.e., miRNA-like RNAs with lengths ranging from 18 to 38 nt.

Importantly, our novel methodology carries several advantages over existing computational methods:

1. Compared to current, purely string comparison methods, DEVs take significantly less time to obtain.
2. Better visualization of ncRNAs processing.
3. SURFR data structures consume very little memory thus allowing real-time calculations.
4. Calculus-based modeling can be directly applied to DEVs to understand ncRNA behavior thus providing a mathematical means to study transcriptomic functionality.
5. Our methodology is highly effective and accurate. To be more specific, our wavelet-based analysis on HS typically identifies ncRNA-derived RNA start and end positions with >=95% identity (within 2 nt) to experimentally validated databases like miRbase as opposed to the state-of-the-art methods based on BAM files such as FlaiMapper, which have been reported to correctly predict 89% of miRNA start positions and 54% of miRNA end positions(78).
6. We have extended our computational methodology to 400+ organisms and all of their sncRNAs without the necessity to change any algorithmic criteria.
7. Our method can address the dynamism associated with transcriptomic analysis using topological interpretation.

### LAGOOn implementation

Similar to SURFR, with LAGOOn, users with no computational background can quickly and easily analyze and compare raw RNA-Seq datasets to comprehensively evaluate lncRNA expressions as well as the potential for lncRNAs to function as sncRNA hosts, miRNA sponges, antisense RNAs, microprotein transcripts, and/or regulators of genomic enhancers. In short, LAGOOn distinguishes itself from existing platforms through offering parallel, real-time expression analysis and functional prediction. Of note, LAGOOn is essentially based on an extended version of MoVaK alignment that similarly employs SVs to perform sequence alignments. In LAGOOn, however, the algorithm was modified during extension in order to trade time and space complexities within the alignment. A detailed explanation regarding these modifications is provided in **Supplemental Information File 3**.

