## Supplementary Information File 1 for "SALTS – <u>S</u>URFR (sncRNA) <u>A</u>nd <u>L</u>AGOOn (lncRNA) <u>T</u>ranscriptomics <u>S</u>uite"

### SALTS – SURFR (sncRNA) And LAGOOn (lncRNA) Transcriptomics Suite: USER MANUAL

**TOOL 1: SURFR**

While an increasing body of evidence suggests specifically excised sncRNA fragments from an array of ncRNAs exist and are functionally relevant, there are currently no Web-based, user-friendly resources that offer comprehensive sncRNA fragment profiling and discovery, functional prediction, and the identification of significant differential expressions among datasets. To address this gap we present SURFR. SURFR refers to our Short Uncharacterized RNA Fragment Recognition tool that identifies all miRNA, snoRNA, and tRNA fragments (as well as fragments from all other ncRNAs annotated in Ensembl) specifically excised in a given transcriptome provided as either a raw user-generated RNA-seq dataset or NCBI SRR file. In addition, SURFR can also compare individual fragment expressions (as well as the expressions of all full length (not fragmented) sncRNAs) among as many as 30 distinct datasets. Of note, SURFR results are stored on the server indefinitely, protected by powerful state-of-the-art cryptographic algorithms, and can be instantly recalled by the user via entering their session key (obtained by clicking "My Session") in the "Get Results" tab on the SURFR home page (**Figure 1A**).

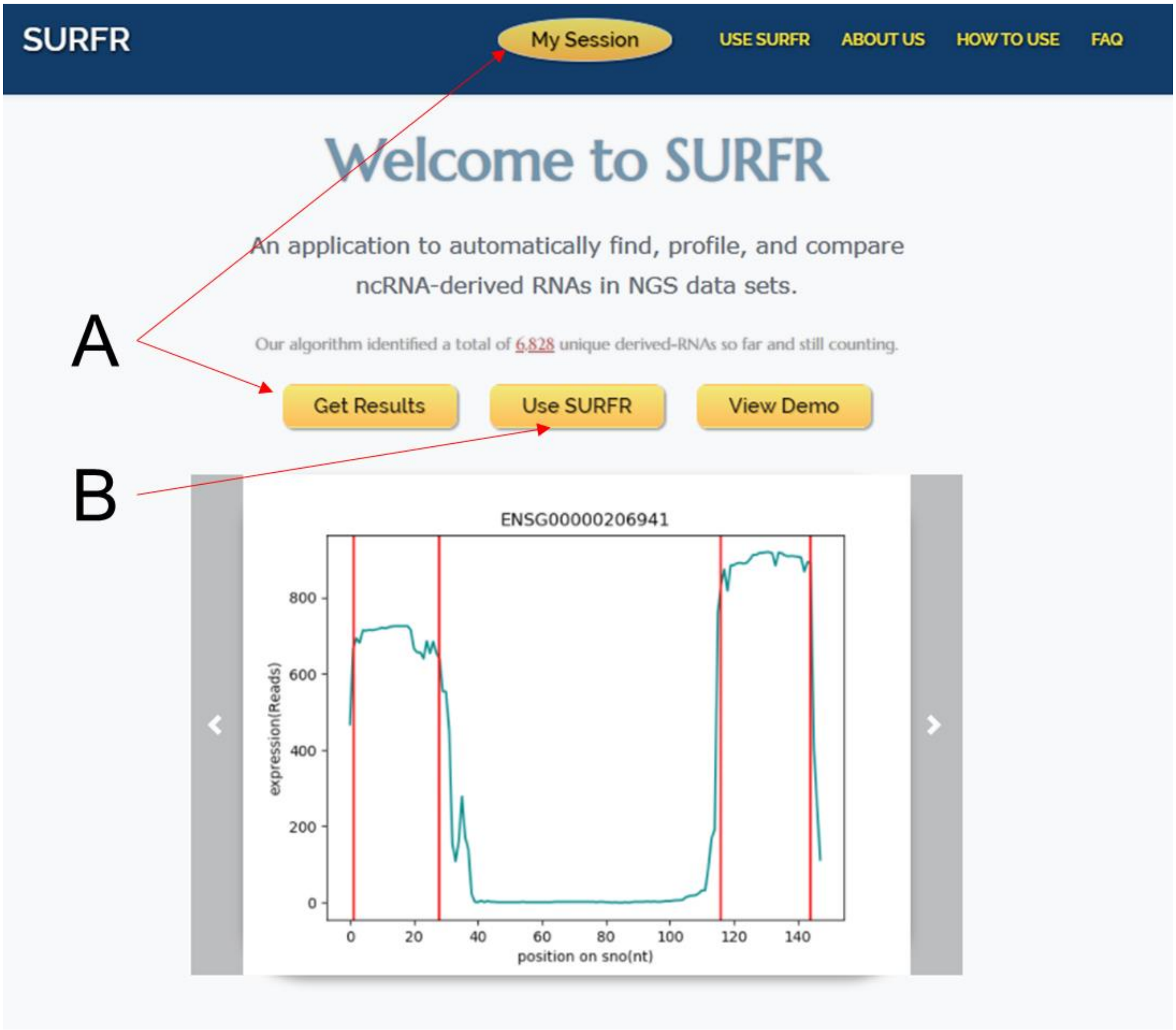


**Figure 1. SURFR home page.** (**A**) My Session and Get Results Links. A single SURFR session can determine ncRNA expressions in up to 10 datasets, and these results can be instantly recalled by the user via entering their session key (obtained by clicking "My Session") in the "Get Results" tab on the SURFR home page. NcRNA expressions in up to 30 datasets can be directly compared by entering multiple session keys in the "Get Results" tab. (**B**) New analyses are performed by following the "Use SURFR" link. "Use SURFR" allows the user to identify all ncRNA fragments and their expressions in up to ten datasets per session.

#### SURFR Input

New analyses are performed by following the "Use SURFR" link (**Figure 1B**). Under "Use SURFR", the user first selects the organism corresponding to the sequences. SURFR small RNA databases have been prepopulated for 440 species including 286 metazoans, 62 plants, and 92 other fungi, protists, and bacteria. Next, the user provides one to ten small RNA sequencing datasets as input. **Figure 2** demonstrates that, these datasets can be all uploaded directly by the user, or all downloaded from the NCBI SRA database(1) by entering SRA IDs (e.g., SRR6495855, SRR4217122), or any combination thereof ( for example, three datasets uploaded by the user along with seven datasets downloaded from the NCBI SRA database). Importantly, a major strength of SURFR is that users can upload most raw small RNA-seq files directly as original, unmodified, compressed FASTQ files (as provided by commercial sequencers) with absolutely no preprocessing and with no specifics about library generation, linkers, or oligonucleotides required. There is no limit on the size of SRA files whereas individual user uploaded files are limited to 1.8 GB regardless of format. As FASTQ compression typically reduces file size by ~70%, individual FASTQ files of up to 6 GB (uncompressed) are allowable. Extremely large sequencing files exceeding even this size can be converted to FASTA format then compressed allowing sequencing files initially > 10 GB to be uploaded. Allowable formats for uploading are uncompressed, standard FASTA or FASTQ files or any major compression of either.

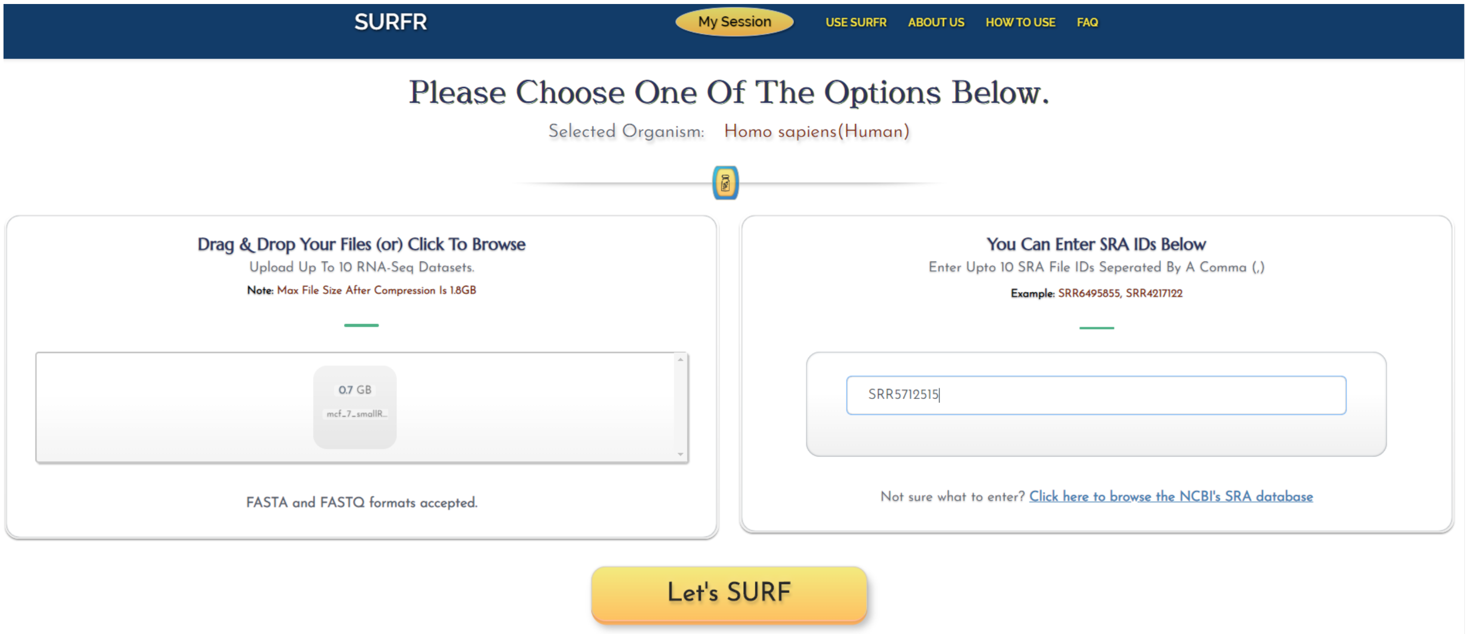


**Figure 2. SURFR input.** SURFR can be used to process and analyze up to ten small RNA sequencing datasets concurrently. The user can input datasets by uploading them directly (left window) and/or specifying publically available datasets to download from the NCBI SRA database by entering SRA IDs (e.g., SRR6495855, SRR4217122) (right window).

#### SURFR Output

After the user uploads/specifies the small RNA-seq datasets and clicks the "Let's SURF" button, the browser is automatically redirected to a report page, progress indicators for each uploaded dataset are provided under the "Click Here To Choose Your File" drop down menu at the top of the page (**Figure 3A**) with individual datasets having completed analysis indicated by a checkmark. Following completion of analysis, results for the individual file selected are then displayed on the report page and organized into several sections (**Figure 3**).

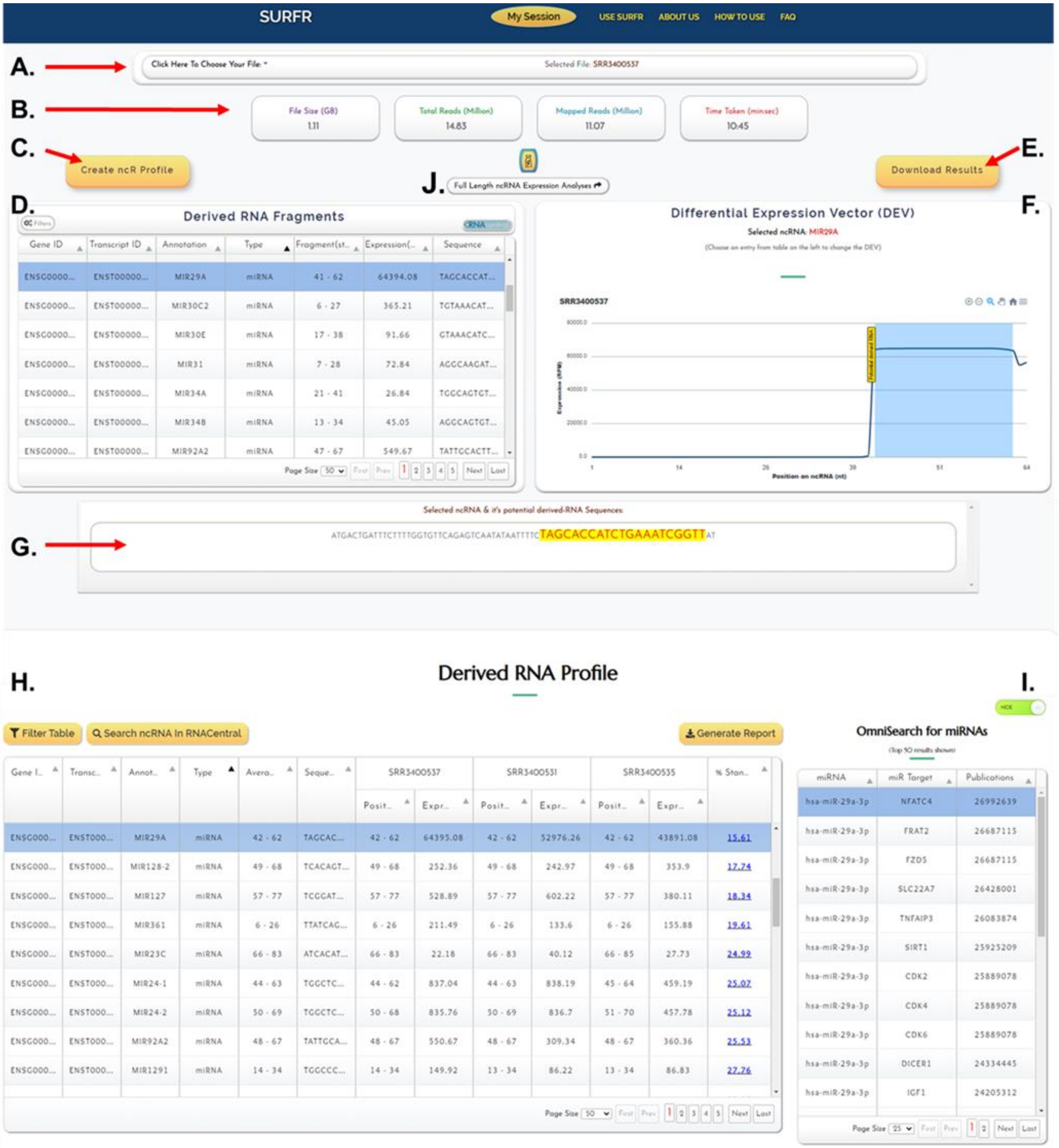


**Figure 3. SURFR report page**. SURFR report example. (**A**) The “Click Here To Choose Your File” drop-down menu for selecting individual RNA-seq files. (**B**) A summary of the overall composition of the selected small RNA-seq dataset. (**C**) The “Create ncR Profile” button automatically populates the derived RNA Profile section at the bottom of the page. (**D**) The “Derived RNA Fragments” window detailing each fragment identified in the individual, selected small RNA-seq dataset. (**E**) The user can download an Excel file detailing the full set of information presented in the “Derived RNA Fragments” window by pressing the “Download Results” button. (**F**) The “Differential Expression Vector (DEV)” window illustrates each nucleotide within a host gene and indicates the fragment called with a blue rectangle. The x-axis represents the position in the ncRNA selected (e.g., miR-29a), and the y-axis depicts the expression levels of the ncRNA at each position. (**G**) The “Selected ncRNA & Called RNA Fragment Sequences” window illustrates the full length host ncRNA (miR-29a) highlighting the SURFR-called fragment in yellow. (**H**) The “Derived RNA Profile” window details each fragment identified in any of the analyzed small RNA-seq datasets and compares fragment expressions across samples. (**I**) The “OmniSearch for miRNAs” window lists the top 50 OmniSearch entries (reported targets and PubMed publications) for an individual miRNA selected in the “Derived RNA Profile” window. (**J**) The “Full Length ncRNA Expression Analyses” button in the upper center of the results page redirects the user to a SURFR window detailing the expressions of all full length sncRNAs in the provided datasets.

A summary of the overall composition of the selected small RNA-seq dataset, including the file size, total number of reads, number of mapped reads, and time taken for analysis is included just below the file selection window at the top of the page (**Figure 3B**).

The user can compare fragment expressions across all datasets by pressing the “Create ncR Profile” button that automatically populates the derived RNA Profile section at the bottom of the page (**Figure 3C**).

The “Derived RNA Fragments” window (**Figure 3D**) details the Ensembl Gene ID, Ensembl Transcript ID, gene annotation (name), the type of gene a fragment was excised from, the start and end positions of a fragment within its host gene, the expression of a fragment in reads per million (RPM), and the nucleotide sequence for each fragment identified in the individual, selected small RNA-seq dataset. The “Derived RNA Fragments” window is an interactive table that allows users to view, sort, and filter small RNA fragments based on any column value. Users can also view host gene information available at the RNAcentral browser by selecting a fragment in the table and then clicking the “RNAcentral” button on the toolbar(2).

The user can download an Excel file detailing the full set of information presented in the “Derived RNA Fragments” window (**Figure 3D**) for each fragment identified in the individual, selected small RNA-seq dataset by pressing the “Download Results” button (**Figure 3E**). An Excel file containing the derived RNA fragment information in its entirety will be automatically downloaded to the user’s computer (**Figure 4**).

| Gene ID | Transcript ID | Annotation | Type | Fragment(start-end) | Expression(RPM) | Sequence |
|---|---|---|---|---|---|---|
| ENSG00000199135.1 | ENST00000362265.1 | MIR101-1 | miRNA | 46 - 66 | 68602 | TACAGTACTGTGATAACTGA |
| ENSG00000284032.1 | ENST00000362111.4 | MIR29A | miRNA | 41 - 62 | 64394 | TAGCACCATCTGAAATCGGTT |
| ENSG00000207752.1 | ENST00000385019.1 | MIR199A1 | miRNA | 46 - 67 | 34071 | ACAGTAGTCTGCACATTGGTT |
| ENSG00000207638.1 | ENST00000384906.1 | MIR99A | miRNA | 12 - 33 | 33760 | AACCCGTAGATCCGATCTTGT |
| ENSG00000288462.1 | ENST00000673161.1 | MIR23A | miRNA | 44 - 63 | 13936 | ATCACATTGCCAGGGATTT |
| ENSG00000198973.4 | ENST00000362103.4 | MIR375 | miRNA | 39 - 60 | 6214 | TTTGTTCGTTCGGCTCGCGTG |
| ENSG00000199085.3 | ENST00000362215.3 | MIR148A | miRNA | 43 - 64 | 3774 | TCAGTGCACTACAGAACTTTG |
| ENSG00000199047.3 | ENST00000362177.3 | MIR378A | miRNA | 42 - 63 | 2166 | ACTGGACTTGGAGTCAGAAGG |
| ENSG00000207713.3 | ENST00000384980.3 | MIR200C | miRNA | 43 - 65 | 1713 | TAATACTGCCGGGTAATGATGG |
| ENSG00000277864.1 | ENST00000516881.1 | SCARNA15 | scaRNA | 65 - 86 | 1360 | AGGTAGATAGAACAGGTCTTG |
| ENSG00000277947.1 | ENST00000619178.1 | SNORD3D | snoRNA | 194 - 217 | 1304 | GGAGAGAACGCGGTCTGAGTGGT |

**Figure 4. Derived RNA Fragments “Download Results” File**. The first few rows of an example “Download Results” Excel file detailing the full set of information presented in the “Derived RNA Fragments” window: Ensembl “Gene ID”, Ensembl “Transcript ID”, gene “Annotation” (name), the “Type” of gene a fragment was excised from, the start and end positions of a fragment within its host gene, the expression of a fragment in reads per million (RPM), and the nucleotide “Sequence” for each fragment identified in the selected small RNA-seq dataset.

| Gene ID | Transcript ID | Annotation | Type | Average start-end | Sequence | SRR3400537 | | SRR3400531 | | SRR3400535 | | % Standard Deviation |
|---|---|---|---|---|---|---|---|---|---|---|---|---|
| | | | | | | Position | Expression | Position | Expression | Position | Expression | |
| ENSG00000199135.1 | ENST00000362265.1 | MIR101-1 | miRNA | 47 – 66 | TACAGTACTGTGATAACTGA | 47 - 66 | 68603 | 47 - 66 | 52890 | 47 - 66 | 57029 | 11.18 |
| ENSG00000284032.1 | ENST00000362111.4 | MIR29A | miRNA | 42 - 62 | TAGCACCATCTGAAATCGGTT | 42 - 62 | 64395 | 42 - 62 | 52976 | 42 - 62 | 43891 | 15.61 |
| ENSG00000207752.1 | ENST00000385019.1 | MIR199A1 | miRNA | 47 - 67 | ACAGTAGTCTGCACATTGGTT | 47 - 67 | 34072 | 47 - 67 | 40080 | 47 - 67 | 36707 | 6.65 |
| ENSG00000207638.1 | ENST00000384906.1 | MIR99A | miRNA | 13 - 33 | AACCCGTAGATCCGATCTTGT | 13 - 33 | 33761 | 13 - 33 | 82093 | 13 - 33 | 60709 | 33.6 |
| ENSG00000288462.1 | ENST00000673161.1 | MIR23A | miRNA | 45 - 63 | ATCACATTGCCAGGGATTT | 45 - 63 | 13937 | 45 - 63 | 8318 | 45 - 64 | 6099 | 34.9 |
| ENSG00000198973.4 | ENST00000362103.4 | MIR375 | miRNA | 40 - 60 | TTTGTTCGTTCGGCTCGCGTG | 40 - 60 | 6215 | 40 - 60 | 1862 | 40 - 60 | 3306 | 47.72 |
| ENSG00000199085.3 | ENST00000362215.3 | MIR148A | miRNA | 44 - 64 | TCAGTGCACTACAGAACTTTG | 44 - 64 | 3775 | 44 - 64 | 4515 | 44 - 64 | 7475 | 30.42 |
| ENSG00000199047.3 | ENST00000362177.3 | MIR378A | miRNA | 43 - 63 | ACTGGACTTGGAGTCAGAAGG | 43 - 63 | 2167 | 43 - 63 | 4704 | 43 - 63 | 3280 | 30.68 |
| ENSG00000207713.3 | ENST00000384980.3 | MIR200C | miRNA | 44 - 65 | TAATACTGCCGGGTAATGATGG | 44 - 65 | 1714 | 44 - 65 | 2322 | 44 - 65 | 7614 | 68.23 |
| ENSG00000277864.1 | ENST00000516881.1 | SCARNA15 | scaRNA | 66 - 86 | AGGTAGATAGAACAGGTCTTG | 66 - 86 | 1361 | 66 - 86 | 843 | 66 - 86 | 958 | 21.05 |
| ENSG00000263934.4 | ENST00000620232.1 | SNORD3A | snoRNA | 196 - 217 | GAGAGAACGCGGTCTGAGTGGT | 195 - 217 | 1305 | 196 - 217 | 294 | 195 - 217 | 921 | 49.61 |
| ENSG00000264940.4 | ENST00000620446.1 | SNORD3C | snoRNA | 196 - 217 | GAGAGAACGCGGTCTGAGTGGT | 195 - 217 | 1305 | 196 - 217 | 294 | 195 - 217 | 921 | 49.61 |
| ENSG00000212409.1 | ENST00000391107.1 | RNY4P18 | misc_RNA | 62 - 86 | CCCCCACTGCTAAATTTGACTGGCT | 62 - 86 | 1014 | 62 - 86 | 360 | 62 - 86 | 235 | 63.76 |
| ENSG00000199172.3 | ENST00000362302.3 | MIR331 | miRNA | 61 - 80 | GCCCCTGGGCCTATCCTAGA | 61 - 80 | 1013 | 0 - 0 | 1 | 0 - 0 | 1 | 141 |
| ENSG00000238711.1 | ENST00000459254.1 | RNY4P25 | misc_RNA | 69 - 93 | CCCCCACTGCTAAATTTGACTGGCT | 69 - 93 | 1013 | 69 - 93 | 360 | 69 - 93 | 234 | 63.73 |
| ENSG00000199017.2 | ENST00000362147.2 | MIR1-1 | miRNA | 46 - 64 | TGGAATGTAAAGAAGTATG | 46 - 65 | 931 | 46 - 65 | 2095 | 46 - 65 | 1026 | 39.07 |

**Figure 5. Derived RNA Profile “Generate Report” File**. The first few rows of an example “**Generate Report**” Excel file detailing the full set of information presented in the “Derived RNA Profile” window.

The “OmniSearch for miRNAs” window (**Figure 3I**) returns the top 50 OmniSearch entries(4) (reported targets and PubMed entries) for an individual miRNA selected in the preceding “Derived RNA Profile” window.

And finally, when desired, the “Full Length ncRNA Expression Analyses” button (**Figure 3J**) redirects the user to a SURFR window detailing the expressions of all full length sncRNAs in the provided datasets regardless of fragmentation. Importantly, all pertinent features (e.g. expression table downloads) described above are similarly available for full length sncRNA analyses via this resource.

#### TOOL 2: LAGOOn

Importantly, LAGOOn is the first Web-based, user-friendly resource that offers real-time lncRNA profiling, the identification of significant differential expressions among datasets, and an array of functional prediction assessments beyond standard mRNA interaction characterizations. Full details of these novel computational methodologies are described in length in **Supplemental Information File 2**.

##### LAGOOn Workflow

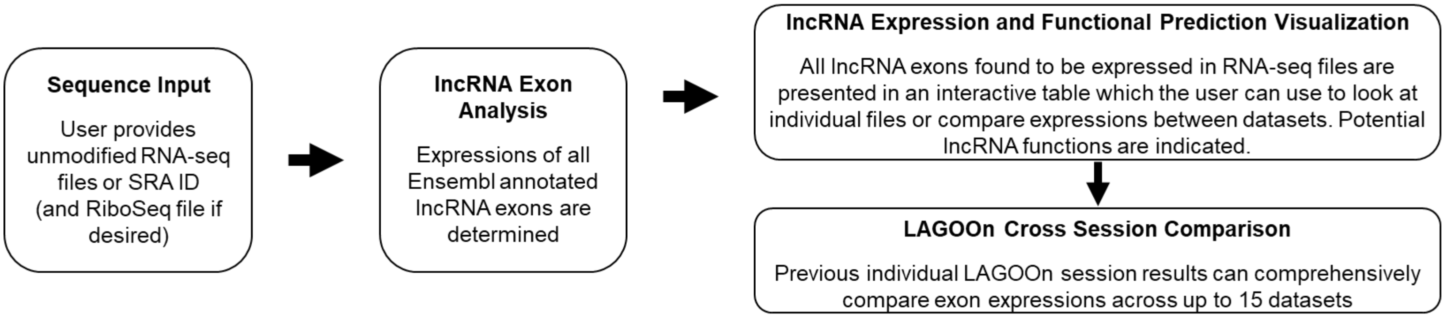


**Figure 7. LAGOOn workflow.** Sequence Input (left). The user provides up to two unmodified RNA-seq files and one Ribo-seq dataset (optional) as input. These datasets can all be uploaded directly by the user or downloaded from the NCBI SRA database by entering SRA IDs. lncRNA Exon Analysis (middle). LAGOOn enumerates all annotated lncRNA expressions in up to three datasets per session. lncRNA Expression and Functional Prediction Visualization (top right). An interactive table is generated comparing the expressions of all exons within individual datasets and comparing exon expressions across all datasets. Tables indicating putative lncRNA functions are also depicted. LAGOOn Cross Section Comparison (bottom right). The user can comprehensively compare all exon expressions identified in up to 15 individual datasets by entering multiple LAGOOn session IDs from separate analyses.

##### LAGOOn Input

As summarized in **Figure 7**, after selecting “Start New Analysis” on the LAGOOn homepage (**Figure 8**), the browser is redirected to the “Data Transfer Options” page where the user provides one or two RNA sequencing datasets as input and is given the chance to provide an optional, additional input, i.e., a Ribo-seq dataset for determining microprotein coding potentials. These datasets can all be uploaded directly by the user, or all downloaded from the NCBI SRA database(1) by entering SRA IDs (e.g., SRR9729388, SRR6290085), or any combination thereof. Importantly, a major strength of LAGOOn is that users can upload most raw RNA-seq files directly as original, unmodified, compressed FASTQ files (as provided by commercial sequencers) with absolutely no preprocessing and with no specifics about library generation, linkers, or oligonucleotides required. There is no limit on the size of SRA files whereas individual user uploaded files are limited to 18 GB regardless of format meaning extremely large sequencing files exceeding even this size can be converted to FASTA format then compressed prior to being uploaded if necessary. Allowable uploaded formats are uncompressed, standard FASTA or FASTQ files or any major compression of either. In addition to this, the LAGOOn homepage provides links to: (1) “Access Your Results” where users can retrieve results from previous sessions via providing a session key and then compare results from up to five separate sessions. (2) “LAGOOn Search” where users can obtain detailed, comprehensive functional predictions for individual lncRNAs. And, (3) “Download Our Databases” where users can download databases containing all the lncRNAs and/or lncRNA exons employed by LAGOOn (**Figure 8**).

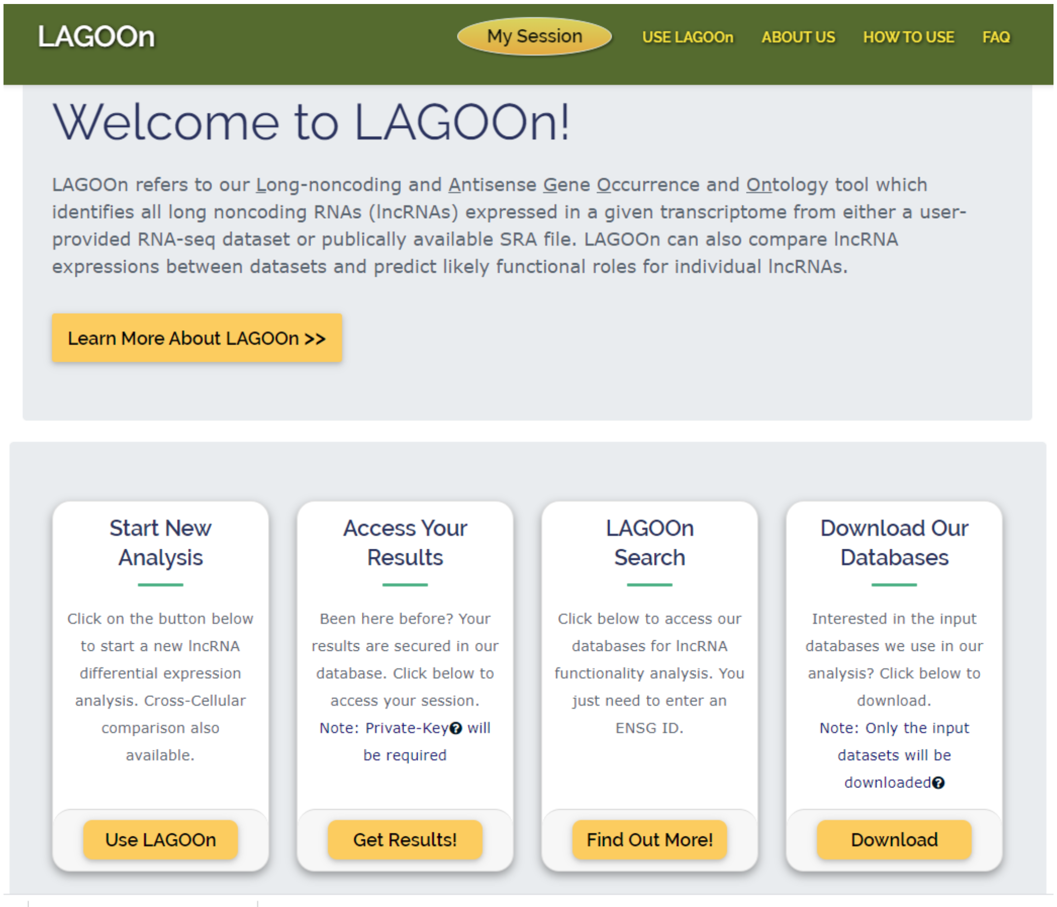


**Figure 8. LAGOOn input.** LAGOOn can be used to process and analyze two RNA-seq datasets as well as one Ribo-seq dataset concurrently. Once the "Start New Analysis" link is selected, the user can input datasets by uploading them directly and/or specifying publically available datasets to download from the NCBI SRA database by entering SRA IDs by selecting the "Start New Analysis" link (bottom left). Users can also retrieve results from previous sessions through providing a session key and compare results from up to five separate sessions by selecting the "Access Your Results" link (bottom middle left). (2) Users can obtain detailed, comprehensive functional predictions for individual lncRNAs by selecting the "LAGOOn Search" link (bottom middle right). And, users can download databases containing all the lncRNAs and/or lncRNA exons employed by LAGOOn by selecting the "Download Our Databases" link (bottom right).

#### LAGOOn Output

After the user uploads/specifies the RNA-seq datasets, the browser is automatically redirected to the LAGOOn report page (**Figure 9**). Initially, a summary of the size and composition of individual RNA-seq datasets, the number of lncRNAs expressed in a dataset, and the top ten most highly expressed lncRNAs in the specified dataset are shown. Following selection of either one or all of the RNA-seq files and the Ribo-Seq file (if included) analyzed from the file selection toolbar (**Figure 9A**), results for the file(s) selected are then displayed on the report page under the "Results" tab (**Figure 9B**), and organized into several distinct sections.

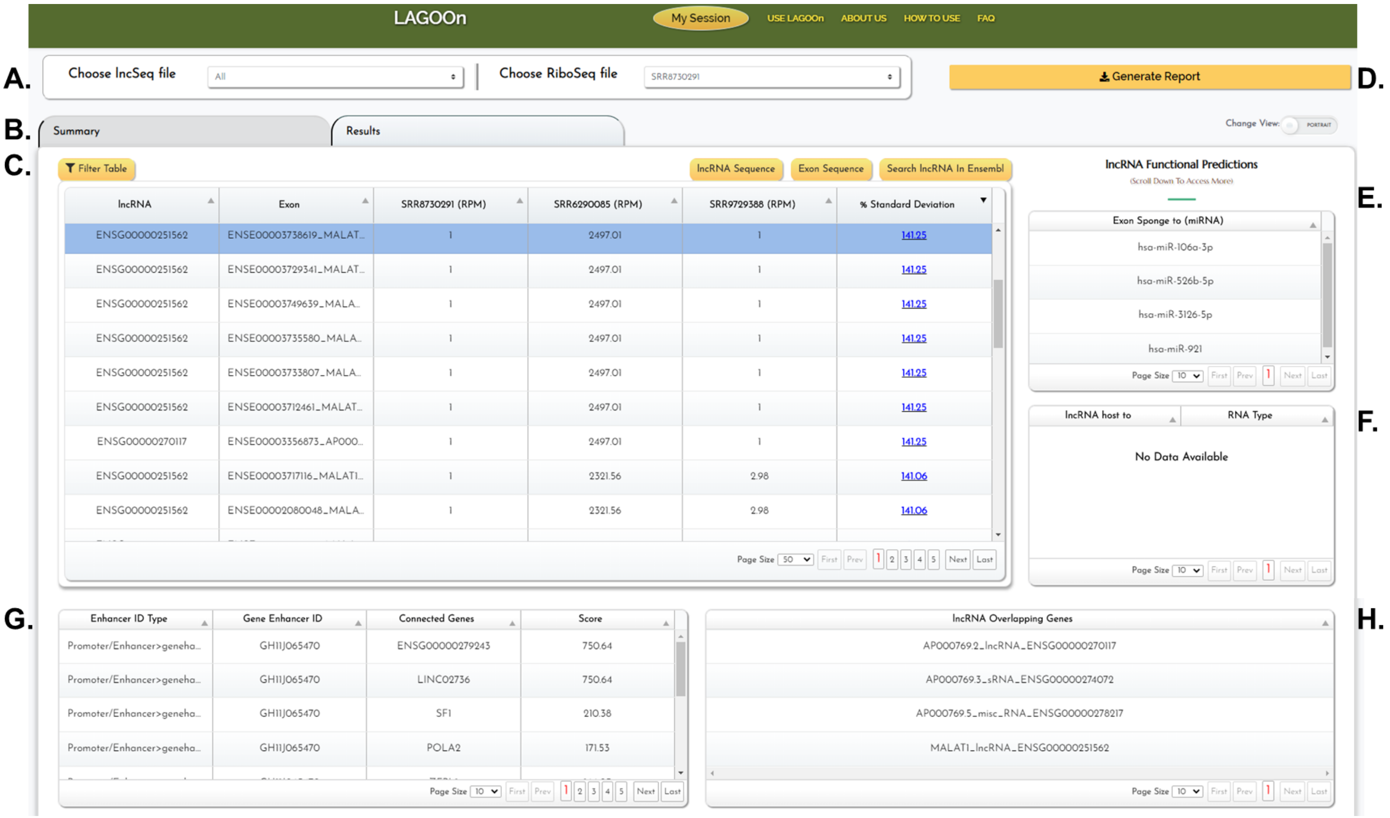


<u>**Figure 9. LAGOOn report page**. LAGOOn report example. (**A**) The file selection toolbar contains drop-down menus for selecting individual RNA-seq and Ribo-Seq files. (**B**) The toolbar allowing selection of either the "Summary" or "Results" tab. (**C**) The lncRNA expression window displays a filterable table of all lncRNA exons expressed in any of the user-provided files. Full length lncRNA sequence, individual exon sequence, or Ensembl lncRNA gene information is obtained by selecting an exon in the table and then clicking the "lncRNA Sequence," "Exon Sequence," or "Search lncRNA in Ensembl" button on the toolbar. (**D**) The "Generate Report" button creates and automatically downloads an Excel file detailing the full set of information presented in the expression table window. (**E**) The "Exon Sponge to (miRNA)" window lists all miRNA complementarities of ten base pairs or greater occurring within the selected lncRNA exon (**F**) The "lncRNA host to" window lists all full length ncRNAs contained in any of the selected lncRNA's exons. (**G**) The "Enhancer" window lists all overlaps between a selected lncRNA and GeneHancer annotated enhancer (as well as genes with expression linked to individual enhancers). (**H**) The "lncRNA Overlapping Genes" window lists all genes even partially overlapping a lncRNA locus on either strand.</u>

The table presented in **Figure 9C** details the Ensembl Gene ID, Ensembl Exon ID along with gene annotation (name), and expressions (RPM) of all lncRNA exons in each individual RNA-seq dataset, and finally, the % standard deviation of the expression of individual exons(3). Importantly, the full list of all exons found to be expressed in any of the datasets is presented. In addition, the expression table is interactive and allows user to view, sort, and filter based on any column value by clicking the "Filter Table" button on the toolbar. Users can also obtain a full length lncRNA sequence, a specific exon sequence, or view the lncRNA gene information available at Ensembl by selecting an exon in the table and then clicking the "lncRNA Sequence," "Exon Sequence," or "Search lncRNA in Ensembl" button on the toolbar.

The user can also download an Excel file detailing the full set of information presented in the expression table window by pressing the "Generate Report" button at the top right of the window (**Figure 9D**). An Excel file containing the expression table window information in its entirety will be automatically downloaded to the user's computer (**Figure 10**). In addition, refined Excel file reports can be downloaded following the application of specific filters (e.g., lncRNAs with RPM > 1 in the Ribo-Seq dataset).

| lncRNA | Exon | SRR8730291 (RPM) | SRR6290085 (RPM) | SRR9729388 (RPM) | % Standard Deviation |
|---|---|---|---|---|---|
| ENSG00000230590 | ENSE00003874886_FTX_-1_FTX transcript, XIST regulator [HGNC:37190]_lncRNA | 1 | 128 | 30 | 102.23 |
| ENSG00000230590 | ENSE00003858311_FTX_-1_FTX transcript, XIST regulator [HGNC:37190]_lncRNA | 1 | 128 | 30 | 102.23 |
| ENSG00000230590 | ENSE00003847528_FTX_-1_FTX transcript, XIST regulator [HGNC:37190]_lncRNA | 1 | 128 | 30 | 102.23 |
| ENSG00000225470 | ENSE00003808225_JPX_1_JPX transcript, XIST activator [HGNC:37191]_lncRNA | 1 | 128 | 30 | 102.23 |
| ENSG00000230590 | ENSE00003241026_FTX_-1_FTX transcript, XIST regulator [HGNC:37190]_lncRNA | 1 | 128 | 30 | 102.23 |
| ENSG00000230590 | ENSE00003429313_FTX_-1_FTX transcript, XIST regulator [HGNC:37190]_lncRNA | 1 | 128 | 30 | 102.23 |
| ENSG00000230590 | ENSE00003861803_FTX_-1_FTX transcript, XIST regulator [HGNC:37190]_lncRNA | 1 | 128 | 30 | 102.23 |
| ENSG00000230590 | ENSE00003849720_FTX_-1_FTX transcript, XIST regulator [HGNC:37190]_lncRNA | 1 | 128 | 30 | 102.23 |
| ENSG00000283117 | ENSE00003789008_AC004949.1_-1_novel transcript_lncRNA | 1 | 35 | 19 | 76.18 |
| ENSG00000284722 | ENSE00003811861_AP003175.1_-1_novel transcript_lncRNA | 1 | 118 | 14 | 118.23 |
| ENSG00000259234 | ENSE00002540221_ANKRD34C-AS1_-1_ANKRD34C antisense RNA 1 [HGNC:48618]_lncRNA | 1 | 54 | 10 | 106.57 |
| ENSG00000259234 | ENSE00002554868_ANKRD34C-AS1_-1_ANKRD34C antisense RNA 1 [HGNC:48618]_lncRNA | 1 | 54 | 10 | 106.57 |
| ENSG00000259234 | ENSE00002573893_ANKRD34C-AS1_-1_ANKRD34C antisense RNA 1 [HGNC:48618]_lncRNA | 1 | 54 | 10 | 106.57 |
| ENSG00000259234 | ENSE00002557951_ANKRD34C-AS1_-1_ANKRD34C antisense RNA 1 [HGNC:48618]_lncRNA | 1 | 54 | 10 | 106.57 |
| ENSG00000259234 | ENSE00002541714_ANKRD34C-AS1_-1_ANKRD34C antisense RNA 1 [HGNC:48618]_lncRNA | 1 | 54 | 10 | 106.57 |
| ENSG00000213904 | ENSE00003224994_LIPE-AS1_1_LIPE antisense RNA 1 [HGNC:48589]_lncRNA | 1 | 17 | 8 | 75.31 |
| ENSG00000213904 | ENSE00003062809_LIPE-AS1_1_LIPE antisense RNA 1 [HGNC:48589]_lncRNA | 1 | 17 | 8 | 75.31 |
| ENSG00000213904 | ENSE00001552276_LIPE-AS1_1_LIPE antisense RNA 1 [HGNC:48589]_lncRNA | 1 | 17 | 8 | 75.31 |
| ENSG00000213904 | ENSE00002995358_LIPE-AS1_1_LIPE antisense RNA 1 [HGNC:48589]_lncRNA | 1 | 17 | 8 | 75.31 |
| ENSG00000251259 | ENSE00002021304_AC004069.1_-1_novel transcript_lncRNA | 1 | 65 | 6 | 120.52 |
| ENSG00000272430 | ENSE00003695861_LINC02637_1_long intergenic non-protein coding RNA 2637 [HGNC:54120]_lncRNA | 1 | 9 | 5 | 65.52 |
| ENSG00000237491 | ENSE00002920037_AL669831.5_1_novel transcript_lncRNA | 1 | 4 | 3 | 47.86 |
| ENSG00000237491 | ENSE00001642276_AL669831.5_1_novel transcript_lncRNA | 1 | 4 | 3 | 47.86 |
| ENSG00000237491 | ENSE00001741526_AL669831.5_1_novel transcript_lncRNA | 1 | 4 | 3 | 47.86 |
| ENSG00000251562 | ENSE00003717116_MALAT1_1_metastasis associated lung adenocarcinoma transcript 1 [HGNC:29665]_lncRNA | 1 | 2322 | 3 | 141.06 |
| ENSG00000251562 | ENSE00002080048_MALAT1_1_metastasis associated lung adenocarcinoma transcript 1 [HGNC:29665]_lncRNA | 1 | 2322 | 3 | 141.06 |
| ENSG00000251562 | ENSE00003742980_MALAT1_1_metastasis associated lung adenocarcinoma transcript 1 [HGNC:29665]_lncRNA | 1 | 2322 | 3 | 141.06 |
| ENSG00000251562 | ENSE00003753954_MALAT1_1_metastasis associated lung adenocarcinoma transcript 1 [HGNC:29665]_lncRNA | 1 | 2146 | 3 | 141.03 |

**Figure 10. lncRNA expression table "Generate Report" File**. The first few rows of an example "**Generate Report**" Excel file detailing the full set of information presented in the lncRNA expression window.

Finally, putative functional roles for lncRNAs/lncRNA exons selected in the expression table are depicted in **Figure 9E-H**. As lncRNAs frequently function as miRNA sponges that directly basepair with and effectively inactivate mature miRNAs(5), the "Exon Sponge to (miRNA)" window lists all miRNA complementarities of ten base pairs or greater occurring within the selected lncRNA exon (**Figure 9E**). Next, as numerous lncRNAs have been shown to encode sncRNAs (e.g., miRNAs and snoRNAs) in their exonic sequences, and sncRNA expression often relies on excision from the host lncRNA transcript(6), the "lncRNA host to" window lists all full length ncRNAs contained any of the selected lncRNA's exons (**Figure 9F**). In addition, as several lncRNAs have been reported to function through regulating the accessibility of transcriptional enhancers overlapping their genomic loci(7), all overlaps between a selected lncRNA and GeneHancer(8) annotated enhancer (and genes with expression linked to individual enhancers) are detailed in the "Enhancer" window (**Figure 9G**). And finally, in addition to lncRNA exonic sequences serving as sncRNA hosts, many sncRNAs are processed from lncRNA introns(6). Furthermore, many lncRNAs serve as naturally occurring antisense silencers of genes located on the strand opposite to themselves(9). For both of these reasons, as well as other potential regulatory relationships, all genes overlapping a lncRNA locus on either the positive or negative strand are detailed in the "lncRNA Overlapping Genes" window (**Figure 9H**). Importantly, a comprehensive report detailing each of the functional predictions is also available for individual lncRNAs by selecting the "LAGOOn Search" button on the homepage after entering a lncRNA Ensembl gene identifier. Notably, this search functionality does not require full LAGOOn analysis.
