## Supplementary Information File 2 for "SALTS – <u>S</u>URFR (sncRNA) <u>A</u>nd <u>L</u>AGOOn (lncRNA) <u>T</u>ranscriptomics <u>S</u>uite"

### Supplemental Information File 2. SURFR: Algorithm for the identification and analysis of ncRNA-derived RNAs

*Abstract*— **Noncoding RNAs (ncRNAs) regulate gene expression in essential cellular processes and play key roles in many human diseases. Small nucleolar RNAs (snoRNAs) are a relatively large group of ncRNAs classically thought to function primarily through regulating the ribosome. Importantly, several studies have now identified significant alterations of different snoRNAs in prostate, breast, and lung malignancies and that many of these snoRNAs have been shown to be processed into microRNA-like molecules known as snoRNA-derived RNAs (sdRNAs). Similarly, several small RNAs have also recently been found to be excised from well characterized tRNAs and also suggested to function in both normal cellular metabolism and disease. In this paper, we present a new computational methodology for the identification, analysis, and visualization of ncRNA-derived RNAs (ndRNAs) with linear time and space complexity (named SURFR for Short Uncharacterized RNA Fragment Recognizer). Our algorithm analyses a next-generation sequencing (NGS) file input then directly outputs a list of potential ndRNAs. In this paper, we present two new concepts, one in the field of sequence alignment and the other in the field of derived RNA analysis. In this study, we (1) describe our algorithm and its application, (2) thoroughly explain how to interpret the results of our method, and (3) describe a way to automatically locate the start and end positions of a ndRNA using wavelets. We also provide the results of our algorithm's running time by analyzing several publicly available datasets and finally discuss future research directions to our work.**

***Keywords—derived RNAs, NGS data analysis, microRNA, snoRNA derived RNA, tRNA Fragment, Algorithm***

#### I. Introduction

During the latter half of the 20th century, one of the greatest achievements in genetic research was the meticulous cataloging of epistatic relationships between genetic loci [1]. While new relationships brought new insights, they also created massive networks of seemingly endlessly interacting genetic pathways. In 1993, however, Lee et al. described a new short noncoding RNA that, despite its size, was eventually recognized as an important player in deciphering complex genetic interactions [2]. These small microRNAs (miRs) are only ~20 nucleotides (nt) in length, but are capable of coordinating the expressions of networks of messenger RNAs (mRNAs) through complementary base pairing [2]. Strikingly, over 2,500 unique human miRs (along with over 30,000 miRs across species) have been cloned [3] since the first human miRs were described in 2001 [4-6]. MiR research has now become an area of intense investigation largely due to the ability of a miR to coordinate the expressions of dozens of genes and the realization that miR mis regulations are commonly associated with oncogenesis (reviewed in [7]). Notably, new Next Generation Sequencing (NGS) technologies aimed at evaluating the microRNA transcriptome have unexpectedly revealed the existence of novel RNA fragments derived from other types of small noncoding RNAs [8]. Although initially discarded as nonspecific RNA degradation products, significant evidence now suggests that RNA fragments derived from small nucleolar RNAs (sdRNAs) and transfer RNAs (tRFs) are not simply random degradation products but are instead specifically excised, functional noncoding RNAs [9]. While little is known about the specific functions of many of these recently identified small non-coding RNAs (sdRNAs, and tRFs), they are all generally believed to function similarly to microRNAs regulating gene expression by binding to complementary target sequences [10-12]. Excitingly, over 1,000 distinct sdRNAs have now been reported from various examinations of the small RNA transcriptomes of several species (human, mouse, chicken, Drosophila, Arabidopsis, wheat, yeast, as well as others. [12-15]). Notably, we (and others [12]) find sdRNA processing to be highly specific and frequently conserved across related species.

Only a small fraction of non-coding RNA derived RNAs (ndRNAs) and their functions have been defined to date. Therefore, there is a broad need for fast, reliable, and explicit computational methods to analyze the vast body of available NGS files capable of performing at a rate proportional to continued data creation. As the first step to almost all NGS data analysis methods, sequence alignment plays the most prominent role in the fields of bioinformatics, computational biology, and computational genomics. In this paper, we present an alignment algorithm that is specifically designed for analyzing and identifying ndRNAs.

Defining the identity and prevalence of ndRNAs in NGS data begins with aligning/mapping the reads from an NGS file to a reference ncRNA sequence database/library or to a reference genome typically through using existing alignment software, which generally involve significant time delays. After that, results are analyzed to identify ndRNAs either manually using excel or by using one of the few available tools [16-24]. Most such tools [16-18,21,24] depend on existing alignment software. One such tool is Bowtie [25], which is one of the fastest and most utilized tools for mapping reads to a genome. Another popular tool is NCBI's BLAST [26], which is the most famous tool for local pairwise sequence alignment. Other tools include natively implemented versions of existing alignment algorithms such as Smith-Waterman (SW) [27], Needleman-Wunsch (NW) [28], and Burrows Wheeler Transform (BWT) [29].

Each of the existing tools requires a significant amount of computational expertise limiting their broad utilization. Importantly, almost all of these tools either require the user to choose from different aligners or expect the users to input, aligned results in file formats such as BAM, or a preprocessed format requiring Linux or terminal-based execution [19-21,23]. That said, there are many computational approaches specific to miRNAs which employ machine-learning based, non-comparative methods, target-centered methods, among others[30] but these are not readily adapted to ndRNAs.

To our knowledge, there exists no sequence alignment algorithm specifically designed for identifying ndRNAs. While our method initially focused on ndRNAs processed from known human ncRNAs, we believe it can easily be extended to virtually any other organism. In this article, we (1) describe our algorithm and its application, (2) thoroughly explain how to interpret the results of our method, and (3) describe a way to automatically locate the start and end positions of a ndRNA using wavelets. We also provide the results of our algorithm's running time by analyzing several publicly available datasets and discuss future research directions to our work.

In brief, our sequence aligner and ndRNA identifier (named SURFr for Short Uncharacterized RNA Finder) allows researchers to:

1. Quickly analyze a raw NGS file to determine the expression levels of all currently annotated human ncRNAs through requiring an exact match of at least 18 nt.
2. Visually identify the portions of an ncRNA that are excised and specifically determine the expressions of excised fragments.
3. Accurately decide the start and end positions of the ndRNA within the ncRNA.
4. Verify if any pieces of a ncRNA are differentially expressed between different NGS files.

#### II. Methodology

Our primary objective is to generate a list of ncRNA-derived RNAs from an input NGS file based on the varying expression at each position of the ncRNA. To achieve this, we propose a new, more efficient, sequence alignment algorithm for determining expression. Our method can be divided into two parts: (1) Sequence Alignment and (2) Derived RNA Analysis.

##### A. *Sequence Alignment*

Our alignment methodology consists of five steps: (1) Data collection, (2) K-mer separation, (3) Constructing Aho-Corsick automaton, (4) Similarity Vector (SV) generation, and (5) Differential Expression Vector (DEV) calculation (MoVaK Alignment). Each of the steps is discussed in the following five subsections.

###### *1) Data Collection:*

Up-to-date annotations for miRNA hairpins, snoRNAs and tRNAs were obtained from miRBase [31], a database containing experimentally verified microRNAs, BioMart [32], a centralized database of all Ensembl annotated gene data, and RFam [33,34], a database of ncRNA sequences conserved across species. We include these three types of ncRNAs as each have been confirmed to have functional small RNAs derived from them.

Raw data files are processed to extract the unique identifier (UID) and associated sequence of each ncRNA. The UID can be the 'Ensemble ID' or any ID provided by the dataset. To avoid redundant computations, we identified the RNAs with the same sequence and length and considered only one among such items. We then reduced six or more consecutive characters (A, C, G, or T) to three to exclude simple sequences (e.g. polyadenylation which is the addition of continuous adenine (A) bases to the end of an mRNA). This step is essential as some of these ncRNAs contain continuous characters of 'A' (up to 19) which would inappropriately align with millions of reads from an NGS file with abundant poly-As.

###### *2) K-mer Separation:*

Most RNA-Seq data analysis techniques use long, imperfect sequence matches identified through extension from short initial alignments to calculate differential expressions [35]. In our case, since we are searching for (18-35 nt) sequences derived from larger sequences, initially requiring the identification of exact matches of at least 18 nt long considerably increases efficiency and makes the problem tractable. In an NGS file with ~10-100 million sequences (reads), only a few thousand of the reads will likely match an ncRNA that has an identical 18 consecutive character identity. As such, we created an index with every possible substring of length 18 (18-mer) from all the ncRNA sequences from our database; the UID associated with the ncRNA is used as a reference. Since the number of sequences in our ncRNA dataset is small (less than 4000 sequences), the total number of 18-mers will be very small in terms of memory requirement (~9MB on disk). One of the major issues encountered during index creation is the same 18-mer coming from multiple ncRNAs. To handle this, our index is created in such a way that all the UIDs that have that substring are stored together to avoid having redundant indices.

Along with the 18-mers, we also store the position of the last character of the 18-mer (index starting at 0) from a given sequence. This simple preprocessing step has a huge impact on the overall performance of the algorithm. A sample index of 18-mers looks as follows:

Table 1. 18-mer index with UIDs and positions

| 18-mer | UID | Position in ncRNA |
|---|---|---|
| GGTGTTAGGGCCAACATC | hsa-mir-2909 | 17 |
| GTGTTAGGGCCAACATCT | hsa-mir-2909 | 18 |
| TTGCAATGATGTGAATCT | ENSG00000281907, ENSG00000199744 | 17 |
| CAATCCCGGGTTTCGGCA | GL000250.2/88968-88895 | 73 |

###### *3) Constructing the Aho-Corasick Automaton*

Aligning a large NGS file line by line with all the ncRNA sequences character by character would be highly inefficient. As such, we perform a 'seed and extend' form of alignment, where alignment is only performed when there is at least a seed length match (18 in our case) between the read and a ncRNA. Therefore before proceeding to alignment, we first search a NGS file for all the 18-mers from our index to separate the reads that match to our sequences. To do so, we use the Aho-Corasick (AC) algorithm [36]. AC is an efficient multiple pattern search algorithm with linear search time complexity. That said, before we can use the AC algorithm for searching, our methodology requires heavy preprocessing (constructing an automaton). Notably we address this limitation by preemptive construction of an automaton with all the indices from our 18-mer index and simply reuse this automaton in all subsequent alignments (or until we want to add or delete some indices).

Since the AC algorithm is one of the most famous pattern search algorithms, implementations of it already exist in almost all general-purpose programming languages. In this work, we used python 3.6 (containing several in-built versions of the AC algorithm of which we specifically used pyahocorasick [37]) to construct our final automaton.

*4) Similarity Vector Generation:*

Similarity Vectors (SV) are also key components of our alignment algorithm. Briefly, our method is based on interdependent features: (1) we use four characters to represent the DNA/RNA and as such in every instance there are only four possibilities for a character, and (2) we are searching for patterns for which we already know these possibilities. Any additional characters commonly found in sequencing data such as ‘N’, ‘X’, are translated to the same four characters during the search. Most commonly, these instances (characters other than A, C, G, T) signify low-quality of the read which represents a well-known issue in NGS data analysis. As such, since ‘N’ is seen more often than the other additional symbols, we allow ‘N’ in a read but not any others. i.e., if any read consists of any character other than A, C, T, G or N, we ignore that read because of inherent bad quality. Knowing the exact number of possibilities for each character allows us to introduce so-called ‘Similarity Vectors’ (SV), which can be used to hold all the information about the sequences.

A similarity vector is a binary array representation of the knowledge of each character’s (ACGTN) location in a given string/sequence. Since we are aligning the same known set of RNAs every time, and we exactly know the positions where each symbol occurs , we process each known ncRNA sequence to find the SVs that determine the exact locations of the characters in the sequence. As an example, consider the SV for the following sequence, S: GCCCTCCTGGTGCTTACCACAGGCTGTGTT where ‘m’ represents the length of the sequence, ‘0’ represents a mismatch/false, and ‘1’ represents a match/true. For each character in our alphabet, we create a linear true/false record in reference to every position/location in the sequence to get an SV with respect to that symbol. Since ‘N’ represents A or C or G or T, SV of N contains all 1s.

$S_A$ = [000000000000000100101000000000] (Match S with A)

$S_C$ = [011101100000100011010001000000] (Match S with C)

$S_G$ = [100000001101000000000110010100] (Match S with G)

$S_T$ = [000010010010011000000000101011] (Match S with T)

$S_N$= [111111111111111111111111111111] (Match S with N)

Fig. 1. Similarity Vector for the sequnce ‘S’

From Fig 1, S has the symbol A at positions 16, 18, and 20. Given this, SVs are capable of determining whether that position contains required character or not. Therefore, an SV of the given sequence S, can be represented as SV(S)= {$S_A$, $S_C$, $S_G$, $S_T$,$S_N$}.

*5) MoVaK Alignment:*

Importantly, we are specifically looking for pieces (fragments) of RNAs that are differentially expressed as compared to the remainder of larger RNA molecules (expressed form the same genomic locus). To find such fragments, we present a novel method to represent changes in RNA expression levels at single nucleotide level resolution. The alignment algorithm, called MoVaK, utilizes a combination of automaton, SVs, and ncRNA sequences. The alignment consists of two steps: (1) Keyword Search and (2) DEV Calculation.

*a) Keyword Search:*

The first step is to search the given NGS file for all the keywords using the AC algorithm. Here, ‘Keyword’ means an 18-mer from our index. AC algorithm’s search complexity is not dependent on the number of keywords but on length of the largest keyword whereas its space complexity is directly proportional to the number of keywords. As mentioned, the number of keywords obtained for our ncRNA dataset is only about 9MB.

Every read in the file that matches to a keyword is mapped to the UID associated with the keyword. If we get more than one UID for a read, we map the read to the UID that has the greater number of keywords matched to the read. In the case where more than one UID has the same number of keywords matched, we assign the read to all possible UIDs.

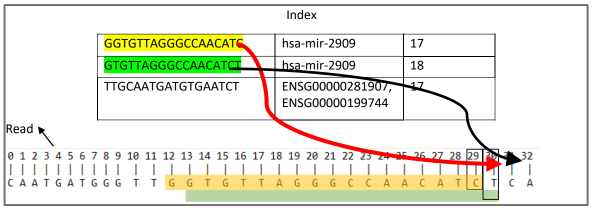

Fig. 2. Representation of matched keywords in a read

For every mapped read, we store the position of each keyword on the read as shown in Fig 2. Since we already have the position of the keyword on the ncRNA, this step should return a list of UIDs each with a set of mapped reads and the position coordinates. As an example, from Fig 2, the position coordinates of the UID, ‘hsa-mir-2909’ for the read would be {(17,29),(18,30)}. After all the reads from a file are scanned and matched reads are assigned to the UIDs, we re-align all the reads that are assigned to a UID.

*b) DEV Calculation:*

To easily identify which specific piece of a given RNA is over expressed, we introduce the concept of DEVs. DEV is an acronym for Differential Expression Vector and is calculated using the SVs. DEVs are 1 dimensional vectors of lengths equal to an ncRNA just like SVs. Unlike SVs, DEVs represent the expression level of the ncRNA at each of its positions. If an ncRNA has no expression in a file, the DEV for that ncRNA is represented as a 1d vector of all 0s . The algorithm for obtaining the DEVs functions as follows:

Obtaining DEV: A DEV for an ncRNA is calculated by pairwise local alignment of all the assigned reads to the reference ncRNA sequence and finally combining the result from all the alignments. Traditionally, in pairwise alignment [27,28], an identity/similarity matrix between the two sequences is constructed and its diagonals are scanned to find the best alignment. In our approach the concept of a matrix is used, but, our method does not actually require constructing one. The algorithm to do this is shown in Fig 4.; we will consider an example to explain the alignment process.

We first start with the traditional matrix examples. Let S= GGCCCTCCTGGTAAA, be an ncRNA sequence with length ‘m’ and the read, R= CTGCATCCACTTGTACTCCTGGAACAATTT with length ‘n’ and for the example’s sake, let’s assume we are looking for an exact match of at least six consecutive characters. Constructing a similarity matrix with matches represented by 1 and mismatches with 0, as shown in Fig 3, needs ‘n*m’ comparisons to populate it and ‘n*m’ space to store the matrix. Then, all the diagonals of the matrix need to be traversed in clockwise direction starting from (0,n) to (m,0) in ‘n+m-1’ (number of diagonals) iterations. Then to check the required consecutive match, each element in every diagonal needs to be verified.

← Read →

← ncRNA →

| | C | T | G | C | A | T | C | C | A | C | T | T | G | T | A | C | T | C | C | T | G | G | A | A | C | A | A | T | T | T |
|---|---|---|---|---|---|---|---|---|---|---|---|---|---|---|---|---|---|---|---|---|---|---|---|---|---|---|---|---|---|---|
| G | 0 | 0 | 1 | 0 | 0 | 0 | 0 | 0 | 0 | 0 | 0 | 0 | 1 | 0 | 0 | 0 | 0 | 0 | 0 | 0 | 1 | 1 | 0 | 0 | 0 | 0 | 0 | 0 | 0 | 0 |
| G | 0 | 0 | 1 | 0 | 0 | 0 | 0 | 0 | 0 | 0 | 0 | 0 | 1 | 0 | 0 | 0 | 0 | 0 | 0 | 0 | 1 | 1 | 0 | 0 | 0 | 0 | 0 | 0 | 0 | 0 |
| C | 1 | 0 | 0 | 1 | 0 | 0 | 1 | 1 | 0 | 1 | 0 | 0 | 0 | 0 | 0 | 1 | 0 | 1 | 1 | 0 | 0 | 0 | 0 | 0 | 1 | 0 | 0 | 0 | 0 | 0 |
| C | 1 | 0 | 0 | 1 | 0 | 0 | 1 | 1 | 0 | 1 | 0 | 0 | 0 | 0 | 0 | 1 | 0 | 1 | 1 | 0 | 0 | 0 | 0 | 0 | 1 | 0 | 0 | 0 | 0 | 0 |
| C | 1 | 0 | 0 | 1 | 0 | 0 | 1 | 1 | 0 | 1 | 0 | 0 | 0 | 0 | 0 | 1 | 0 | 1 | 1 | 0 | 0 | 0 | 0 | 0 | 1 | 0 | 0 | 0 | 0 | 0 |
| T | 0 | 1 | 0 | 0 | 0 | 1 | 0 | 0 | 0 | 0 | 1 | 1 | 0 | 1 | 0 | 0 | 1 | 0 | 0 | 1 | 0 | 0 | 0 | 0 | 0 | 0 | 0 | 1 | 1 | 1 |
| C | 1 | 0 | 0 | 1 | 0 | 0 | 1 | 1 | 0 | 1 | 0 | 0 | 0 | 0 | 0 | 1 | 0 | 1 | 1 | 0 | 0 | 0 | 0 | 0 | 1 | 0 | 0 | 0 | 0 | 0 |
| C | 1 | 0 | 0 | 1 | 0 | 0 | 1 | 1 | 0 | 1 | 0 | 0 | 0 | 0 | 0 | 1 | 0 | 1 | 1 | 0 | 0 | 0 | 0 | 0 | 1 | 0 | 0 | 0 | 0 | 0 |
| T | 0 | 1 | 0 | 0 | 0 | 1 | 0 | 0 | 0 | 0 | 1 | 1 | 0 | 1 | 0 | 0 | 1 | 0 | 0 | 1 | 0 | 0 | 0 | 0 | 0 | 0 | 0 | 1 | 1 | 1 |
| G | 0 | 0 | 1 | 0 | 0 | 0 | 0 | 0 | 0 | 0 | 0 | 0 | 1 | 0 | 0 | 0 | 0 | 0 | 0 | 0 | 1 | 1 | 0 | 0 | 0 | 0 | 0 | 0 | 0 | 0 |
| G | 0 | 0 | 1 | 0 | 0 | 0 | 0 | 0 | 0 | 0 | 0 | 0 | 1 | 0 | 0 | 0 | 0 | 0 | 0 | 0 | 1 | 1 | 0 | 0 | 0 | 0 | 0 | 0 | 0 | 0 |
| T | 0 | 1 | 0 | 0 | 0 | 1 | 0 | 0 | 0 | 0 | 1 | 1 | 0 | 1 | 0 | 0 | 1 | 0 | 0 | 1 | 0 | 0 | 0 | 0 | 0 | 0 | 0 | 1 | 1 | 1 |
| A | 0 | 0 | 0 | 0 | 1 | 0 | 0 | 0 | 1 | 0 | 0 | 0 | 0 | 0 | 1 | 0 | 0 | 0 | 0 | 0 | 0 | 0 | 1 | 1 | 0 | 1 | 1 | 0 | 0 | 0 |
| A | 0 | 0 | 0 | 0 | 1 | 0 | 0 | 0 | 1 | 0 | 0 | 0 | 0 | 0 | 1 | 0 | 0 | 0 | 0 | 0 | 0 | 0 | 1 | 1 | 0 | 1 | 1 | 0 | 0 | 0 |
| A | 0 | 0 | 0 | 0 | 1 | 0 | 0 | 0 | 1 | 0 | 0 | 0 | 0 | 0 | 1 | 0 | 0 | 0 | 0 | 0 | 0 | 0 | 1 | 1 | 0 | 1 | 1 | 0 | 0 | 0 |

Fig. 3. Similarity matrix between S and R highlighting diagonals with at least six consecutive 1s

```
Given the read ‘R’, SV {A, C, T, G, N}, length of read
‘n’, length of ncRNA ‘m’, list of position coordinates,
(position on ncRNA, position on Read) (a_i,
b_i)=[(a_1,b_1),(a_2,b_2),(a_3,b_3)…]
   //Initialize an empty set data structure (holds only
unique values)
   SET E[];
   //Initialize a 1d matrix of length m with 0s.
   Alignment=[0] of length ‘m’
   //repeat for every tuple of position coordinates
   for each (a_i, b_i) {
   //if the difference of ai, bi in E then skip
   if (a_i-b_i) in E:
   continue;
   else:
   //add the value a-b to the set
   add (a_i-b_i) to E;
   j= n-(b_i-a_i)-1;
   k= m-(a_i-b_i)-1;
   start1= min(j+1,m);
   stop1= max(1,j-n+2);
   start2= max(1,k-m+2);
   stop2= min(k+1,n);
   // step = -1 gives output in reverse order
   //we use python’s “arange” function instead of a for
loop here
   //arange(1,4)=[1,2,3,4]     whereas     arange(1,4,-
1)=[4,3,2,1]
   //x= all the x coordinates of the diagonal elements.
   //y= all the y coordinates of the diagonal elements.
   x= for(start1-1; stop1-2; step= -1);
   y= for(start2-1, stop2-2; step= -1);
   //Initialize another 1d matrix to hold diagonal element
values
   Diagonal=array (length m);
   for (int val=0; val < length(x); val++){
   //Fill in the diagonal elements using an SV and
positions (x_j,y_j)
   Diagonal [x[ val ] ]= SV [R [ y [ val ] ] ] [x [ val ]
];
   }
   //Perform 1d matrix addition
   Alignment= Alignment + Diagonal; }
```

Fig. 4. Algorithm to obtain the matched diagonals

For our method, let us assume that the two sequences (ncRNA & read) are given, that we have the SV for the ncRNA as well as a list of position coordinates. We can obtain the diagonals of the similarity matrix along with the required consecutive match, without constructing the matrix or going through all the diagonals; to accomplish this, we use the pseudocode shown in Fig 4.

In their book [38], the authors describe an efficient way to iterate through all the diagonal elements of an nXm matrix using just one for loop and two python’s NumPy arange functions [39]. The arange function, given two integers, produces all the values within the range of the given integers. We use their algorithm to generate coordinates of the diagonal elements and SVs to get values of each element. We use the same example presented earlier, where we are looking for an exact match of at least six consecutive characters, S= GGCCCTCCTGGTAAA, the ncRNA sequence with length ‘m’ and the read, R= CTGCATCCACTTGTACTCCTGGAACAATTT with length ‘n’.

To begin, divide ‘S’ into all possible 6-mers then the index looks as shown in Fig 5. The AC search on R for the keywords (6-mers) would return, [(9,20), (10,21)], and SV(S)={A=000000000000111; C=001110110000000; G=110000000110000; T= 000001001001000; N=111111111111111}

INDEX TABLE FOR S

| 6-mer | UID | Position |
|---|---|---|
| GGCCCT | S | 5 |
| GCCCTC | S | 6 |
| CCCTCC | S | 7 |
| CCTCCT | S | 8 |
| CTCCTG | S | 9 |
| TCCTGG | S | 10 |
| CCTGGT | S | 11 |
| CTGGTA | S | 12 |
| TGGTAA | S | 13 |
| GGTAAA | S | 14 |

Fig. 5. 6-mer index for the example sequence ‘S’

| | C | T | G | C | A | T | C | C | A | C | T | T | G | T | A | C | T | C | C | T | G | G | A | A | C | A | A | T | T | T |
|---|---|---|---|---|---|---|---|---|---|---|---|---|---|---|---|---|---|---|---|---|---|---|---|---|---|---|---|---|---|---|
| G | 0 | 0 | 1 | 0 | 0 | 0 | 0 | 0 | 0 | 0 | 0 | 0 | 1 | 0 | 0 | 0 | 0 | 0 | 0 | 0 | 1 | 1 | 0 | 0 | 0 | 0 | 0 | 0 | 0 | 0 |
| G | 0 | 0 | 1 | 0 | 0 | 0 | 0 | 0 | 0 | 0 | 0 | 0 | 1 | 0 | 0 | 0 | 0 | 0 | 0 | 0 | 1 | 1 | 0 | 0 | 0 | 0 | 0 | 0 | 0 | 0 |
| C | 1 | 0 | 0 | 1 | 0 | 0 | 1 | 1 | 0 | 1 | 0 | 0 | 0 | 0 | 0 | 1 | 0 | 1 | 1 | 0 | 0 | 0 | 0 | 0 | 1 | 0 | 0 | 0 | 0 | 0 |
| C | 1 | 0 | 0 | 1 | 0 | 0 | 1 | 1 | 0 | 1 | 0 | 0 | 0 | 0 | 0 | 1 | 0 | 1 | 1 | 0 | 0 | 0 | 0 | 0 | 1 | 0 | 0 | 0 | 0 | 0 |
| C | 1 | 0 | 0 | 1 | 0 | 0 | 1 | 1 | 0 | 1 | 0 | 0 | 0 | 0 | 0 | 1 | 0 | 1 | 1 | 0 | 0 | 0 | 0 | 0 | 1 | 0 | 0 | 0 | 0 | 0 |
| T | 0 | 1 | 0 | 0 | 0 | 1 | 0 | 0 | 0 | 0 | 1 | 1 | 0 | 1 | 0 | 0 | 1 | 0 | 0 | 1 | 0 | 0 | 0 | 0 | 0 | 0 | 0 | 1 | 1 | 1 |
| C | 1 | 0 | 0 | 1 | 0 | 0 | 1 | 1 | 0 | 1 | 0 | 0 | 0 | 0 | 0 | 1 | 0 | 1 | 1 | 0 | 0 | 0 | 0 | 0 | 1 | 0 | 0 | 0 | 0 | 0 |
| C | 1 | 0 | 0 | 1 | 0 | 0 | 1 | 1 | 0 | 1 | 0 | 0 | 0 | 0 | 0 | 1 | 0 | 1 | 1 | 0 | 0 | 0 | 0 | 0 | 1 | 0 | 0 | 0 | 0 | 0 |
| T | 0 | 1 | 0 | 0 | 0 | 1 | 0 | 0 | 0 | 0 | 1 | 1 | 0 | 1 | 0 | 0 | 1 | 0 | 0 | 1 | 0 | 0 | 0 | 0 | 0 | 0 | 0 | 1 | 1 | 1 |
| G | 0 | 0 | 1 | 0 | 0 | 0 | 0 | 0 | 0 | 0 | 0 | 0 | 1 | 0 | 0 | 0 | 0 | 0 | 0 | 0 | 1 | 1 | 0 | 0 | 0 | 0 | 0 | 0 | 0 | 0 |
| G | 0 | 0 | 1 | 0 | 0 | 0 | 0 | 0 | 0 | 0 | 0 | 0 | 1 | 0 | 0 | 0 | 0 | 0 | 0 | 0 | 1 | 1 | 0 | 0 | 0 | 0 | 0 | 0 | 0 | 0 |
| T | 0 | 1 | 0 | 0 | 0 | 1 | 0 | 0 | 0 | 0 | 1 | 1 | 0 | 1 | 0 | 0 | 1 | 0 | 0 | 1 | 0 | 0 | 0 | 0 | 0 | 0 | 0 | 1 | 1 | 1 |
| A | 0 | 0 | 0 | 0 | 1 | 0 | 0 | 0 | 1 | 0 | 0 | 0 | 0 | 0 | 1 | 0 | 0 | 0 | 0 | 0 | 0 | 0 | 1 | 1 | 0 | 1 | 1 | 0 | 0 | 0 |
| A | 0 | 0 | 0 | 0 | 1 | 0 | 0 | 0 | 1 | 0 | 0 | 0 | 0 | 0 | 1 | 0 | 0 | 0 | 0 | 0 | 0 | 0 | 1 | 1 | 0 | 1 | 1 | 0 | 0 | 0 |
| A | 0 | 0 | 0 | 0 | 1 | 0 | 0 | 0 | 1 | 0 | 0 | 0 | 0 | 0 | 1 | 0 | 0 | 0 | 0 | 0 | 0 | 0 | 1 | 1 | 0 | 1 | 1 | 0 | 0 | 0 |

Fig. 6. Relation between SVs and the similarity matrix

As we can see from the highlighted portion of Fig 6, wherever we see a ‘C’ in the read, we see the same pattern of 0s and 1s repeated through the matrix columns as well. If we look closer, the pattern we see in the ‘C’ columns is also same as the ‘ $S_C$’ from SV(S). Similarly $S_A$, $S_G$, and $S_T$ are repeated at A, G, T’s positions on the matrix respectively. Since we already have the SVs, we can access the matrix elements without constructing it. both for this example as well as the general case. SVs are the building blocks of the similarity matrix and by using SVs for every read, we can eliminate ‘n*m’ comparisons and require only the 5n space rather than the n*m to store the matrix.

For example, if average lengths of ncRNAs and reads are 350, 65 respectively, and let 1000 reads are mapped to the ncRNA, we can avoid 350*65*1000=22.75 million computations. For one entire file, if about 1000 out of 4,000 total ncRNAs has 1000 mapped reads each, then 1,000,000 reads* 350*65*1000=22,750,000,000,000 (22.75 trillion) comparisons are eliminated.

Next, assuming we have SV(S); position coordinates P(S)={(9,20),(10,21)}; n=30; m=15, we can use the alignment process shown in fig 7.

```
SET E[];
Alignment=[000000000000000]//length(m)
//[(9,20),(10,21)]
(9, 20) {
if (9-20) in E: //not true
continue;
else:
add (9-20) to E; //E=[-11]
j= 30-(20-9)-1;//=18
k= 15-(9-20)-1;//=25
start1= min(18+1,15);//=15
stop1= max(1,18-30+2);//=1
start2= max(1,25-15+2);//=12
stop2= min(25+1,30);//26
x= for(15-1;1-2;-1);
//x= [14,13,12,11,10,9,8,7,6,5,4,3,2,1,0]
y= for(12-1,26-2;-1);
//y= [25,24,23,22,21,20,19,18,17,16,15,14,13,12,11]
Diagonal=[000000000000000];
for (int val=0; val < 15; val++){
//val=0
Diagonal [x[ 0 ] ]= SV [R [ y [ 0 ] ] ] [x [ 0 ] ];
// Diagonal [14]=SV[R[25]][14]
// Diagonal [14]=SV[A][14]
// Diagonal [14]=1
// Diagonal =[000000000000001]
//val=1
Diagonal [x[ 1 ] ]= SV [R [ y [ 1 ] ] ] [x [ 1 ] ];
// Diagonal [13]=SV[R[24]][13]
// Diagonal [13]=SV[C][13]
// Diagonal [13]=0
// Diagonal =[000000000000001]
//val=2
Diagonal [12]=1
// Diagonal =[000000000000101]
                              :::
                              :::
Diagonal= [110011111110101]
}
Alignment= [110011111110101];
(10,21) { //not true i.e., (9,20) and (10,21) are on
same diagonal }
```

Fig. 7. Applying SVs for aligning the example sequences

The algorithm shown in Fig 7 produces a 1d integer matrix of length m representing the alignment between the ncRNA sequence and one read. A '0' on the alignment means there is a mismatch between the sequences and '1' means a match at that location i.e., for every alignment, 1 at a position means that position of the RNA is active. The position coordinates play a crucial role in the alignment process by providing us with coordinates of the elements on the required diagonals i.e., because our keyword length is equal to the required minimum match wherever a keyword is matched, those positions on the matrix will contain the minimum required consecutive 1s. So, instead of going through all the diagonals to check for required length match, we can selectively obtain just the required diagonals.

There is a possibility that we can get more than one diagonal with 18 consecutive 1s; we have to consider all such diagonals. To do so, we filter the position coordinates such that we only choose one element from each diagonal, because we only need one element to get entire the diagonal. To determine whether two coordinates (a1,b1) and (a2,b2) lie on the same diagonal, we can check whether (a1-b1)=(a2-b2) or (b1-a1)=(b2-a2). If this is true, then the points lie on the same diagonal else they are from different diagonals.

Every alignment/diagonal obtained from above is added using matrix addition to the DEV of the ncRNA under analysis as shown in Fig 8.

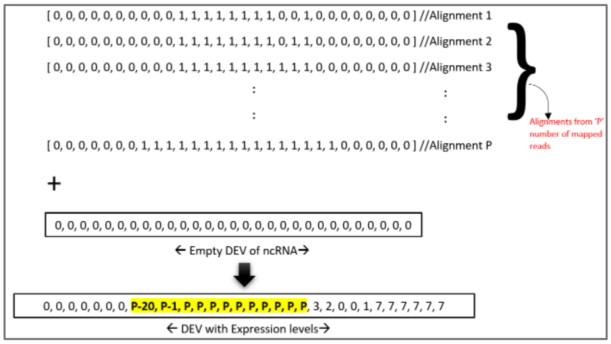

Fig. 8. Calculating the DEV of an ncRNA

We repeat this algorithm for all the mapped reads of the ncRNA sequence. Fig 9 represents a DEV for a ncRNA with a length of 88 characters. The integers in the DEV denote the expression of the ncRNA (reads) at each position. When visualized, the DEV from Fig 9 looks like Fig 10. The x-axis in the graph shown in Fig 10 represents the position on the ncRNA, and the y-axis represents the expression levels of the ncRNA at each position.

```
[924, 953, 955, 954, 945, 958, 954, 959, 959, 962, 962,
964, 964, 961, 965, 964, 964, 964, 957, 951, 941, 928,
928, 925, 895, 917, 907, 908, 693, 813, 628, 798, 616,
793, 844, 537, 33, 14, 14, 15, 15, 15, 17, 18, 20, 19,
101, 161, 208, 863, 1163, 1201, 1343, 1437, 1451, 1477,
1487, 1515, 1522, 1583, 1539, 1945, 3047, 1813, 1934,2105,
10424, 11509, 12593, 12867, 13074, 13073, 13068, 13068,
13070, 13065, 13069, 13068, 13068, 13064, 13061, 13059,
12974, 12916, 12864, 12165, 11770, 11666]
```

Fig. 9. Differential Expression Vector of an ncRNA

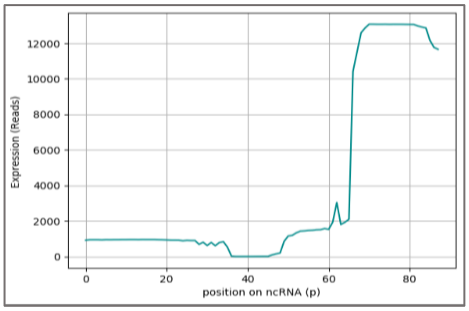

Fig. 10. Visualization of the DEV shown in Fig 9

#### B. *Derived RNA Analysis*

As explained earlier, DEVs are 1 d vectors representing the activity generated by all the positions of a ncRNA. This activity can also be treated as signals generated by each position of the ncRNA within a cell. These signals are mixed with data and noise. The concept of derived RNAs is comparatively new in the field of Biology and Bioinformatics. The derived RNA signals within a longer ncRNA signal are small, clusters in a specific location, surrounded by low expression areas, and vary for each signal. Multiple ndRNAs can be derived from a same ncRNA and each ndRNA's length is independent of the ncRNA. Hence, it is not efficient for us to hardcode the values and limits within the RNA signal. As such we need the flexibility to deal with all of these issues.

In this section, we would like to present a novel means of defining derived RNAs using wavelet analysis on DEVs. A wavelet is a short-lived signal within a continuous longer signal. They are localized in time and frequency and are specific to the particular signal under study [40].

[41] provides a way to detect peaks of a specific range of lengths within a mass spectrometry spectra using continuous wavelet transform (cwt). Peak detection using cwt is sensitive enough to identify multiple side-by-side peaks with different amplitudes. As such we have elected to employ this methodology to automatically detect the presence, expression levels, and start and end positions of novel ndRNAs from our DEVs. We have specifically used Ricker wavelet [42], and python's SciPy [43] 'signal' module to perform our wavelet analysis. Our peak detection algorithm works as follows:

1. Apply continuous wavelet transform on the DEV for the required group of widths (18-35)
2. Find relative maxima for each row connected across adjacent rows in the cwt matrix
3. Finally, based on the noise, filter the relative maxima obtained from step 2 to get the peaks using cwt.

Fig. 11. Algorithm for peak detection on DEVs using cwt

Summary results of our wavelet analyses are discussed in the following section.

#### III. Results and Discussion

Since the concept of DEVs is new, and our primary goal is to align sequencing reads to known ncRNA sequences to efficiently identify derived RNAs, we have compared our results, by aligning the same files (ncRNAs and NGS), using BLAST, which also uses AC algorithm as a base and employs a seed and extend alignment method. BLAST is also one of the fastest and most widely used sequence alignment tools in the bioinformatics community.

All results of our algorithm were obtained using an Intel(R) Core(TM) i7-8550U CPU @1.80GHz 1.99GHz, Windows, laptop computer with 16 GB RAM, 8 cores. Whereas the BLAST results are computed using a machine, with Centos OS, 5-16CPUs, 1GB per core memory, provided by High Performance Computing (HPC), Alabama Supercomputer (ASC) authority [44].

To achieve concurrency in our method, after the keyword search, the mapped reads are assigned to 16 parallel python processes to calculate the DEVs. Similarly, for the BLAST, we used 16 threads in ASC for the performance to be comparable. The only difference is that, BLAST returns local alignment results for the ncRNAs and our method returns DEVs for the ncRNAs. Times taken for both methods to analyze a set of publicly available RNA-Seq data files [45] is provided in Table 2, where, the columns, 'MoVaK time' and 'BLAST time' contain the time taken for analysis of each file using our alignment and BLAST respectively.

Table 2 Comparison of alignment times between MoVaK alignment and NCBI BLAST +

| File ID | Size (GB) | MoVaK time (hr:min:sec) | BLAST time (hr:min:sec) |
|---|---|---|---|
| SRR4451026 | 1.51 | 00:02:39 | 6:13:00 |
| SRR4451007 | 1.77 | 00:05:48 | 5:43:00 |
| SRR4451052 | 1.91 | 00:03:47 | 7:37:00 |
| SRR4451034 | 1.20 | 00:03:13 | 6:57:00 |
| SRR4450996 | 1.43 | 00:04:27 | 6:21:00 |
| SRR4217151 | 0.9 | 00:02:08 | 8:41:00 |
| SRR4217150 | 1.49 | 00:03:02 | 7:58:00 |
| SRR4217130 | 2.72 | 00:05:08 | 8:46:00 |
| DRR036719 | 5.80 | 00:07:15 | 9:18:00 |
| DRR036718 | 6.81 | 00:07:27 | 8:54:00 |
| SRR4451015 | 1.58 | 00:03:31 | 9:06:00 |
| SRR4451037 | 1.75 | 00:06:09 | 5:55:00 |
| SRR4451025 | 1.69 | 00:05:04 | 5:10:00 |
| **AVERAGE** | **2.35** | **00h:04m:35s** | **7h:26m:00** |

On an average, the time necessary for our method to analyze each of 13 different RNA-Seq files was less than five minutes whereas BLAST averaged nearly seven and half hours to analyze each of these same files.

Fig 12 contains four distinct DEV visualizations depicting fragments derived from individual ncRNAs (derived fragment boundaries are represented as vertical red lines). According to our method, these red lines correspond to the start and end positions of the ndRNAs and that the indicated portion of the ncRNA is differentially / more highly expressed than the rest of the full length ncRNA. Fig 12c shows an ncRNA containing two ndRNAs excised from distinct positions. Fig 12d contains no red vertical lines because all of the positions within the ncRNA are similarly prevalent in the aligning sequencing files indicating that no internal regions are specifically excised (differentially expressed) meaning there are no ndRNAs produced by this ncRNA.

To avoid as many false positives as possible, we screen the results obtained from wavelet analysis further so that, only ncRNAs with at least 100 mapped reads are considered, In addition, when there is more than one peak, only peaks that are at least 33.33% of the average expression of the highest peak are considered. Finally, the average of the expressions of all the positions within the start and end positions of the ndRNAs on DEVs is returned as expression of that ndRNA in the given file. Importantly, to evaluate the ability of SURFr to identify actual fragments of noncoding RNAs present within next generation RNA sequencing files we compared the start and end positions of several of our derived fragments with those of previously reported, experimentally verified sequence fragments. Table 3 compares start and end positions of some of the microRNA excised fragments identified in our preliminary SURFr analyses to those of known microRNAs annotated in miRBase. Similarly, we also compared our tRNA fragment calls to those included in MINTbase [46] (a comprehensive, experimentally verified tRNA fragment database), and our snoRNA derived RNA calls to RNA fragments shown to be excised from snoRNAs in specific cell types. Excitingly, we found that all SURFr defined ndRNA start and end positions corresponding to previously annotated, laboratory verified noncoding RNA derived fragments agreed within 2 nucleotides.

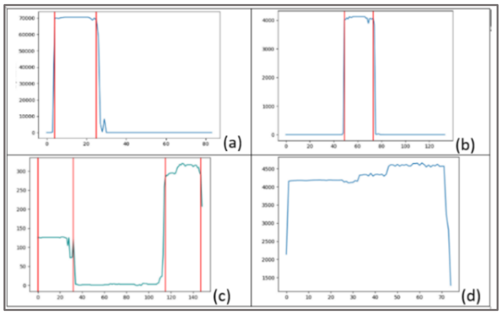

Fig. 12. Results obtained from wavelet analysis on four distinct DEVs

TABLE 3 COMPARISON OF SURFR START AND END POSITION CALLS WITH EXPERIMENTALLY-VERIFIED POSITIONS

| microRNA | SURFr start-end | miRBase start-end |
|---|---|---|
| hsa-mir-148a | 43-64 | 44-65 |
| hsa-mir-101-1 | 46-65 | 47-67 |
| hsa-mir-330 | 57-77 | 57-79 |
| hsa-mir-4446 | 43-62 | 43-64 |
| hsa-mir-152 | 53-73 | 54-74 |
| hsa-mir-127 | 56-76 | 57-78 |
| hsa-mir-361 | 5-25; 43-66 | 6-27; 45-67 |
| hsa-mir-23a | 44-65 | 45-65 |
| hsa-mir-25 | 51-71 | 52-73 |

#### IV. HOW & WHY OUR ALGORITHM WORKS

In this section, we will briefly describe the mathematical theory behind our algorithm. We theorize that, each ncRNA's "expression" is a higher-dimensional function of ACTG throughout its length and is continuously changing based on the state and functionality of the cell. And, a single RNA-seq dataset represents the current physical state of the transcriptome. As such, a complex Hilbert space (HS) in mathematics is a two dimensional inner product space with in infinite dimensional vector space. In quantum mechanics, such Hilbert spaces are used to represent the current state of a physical system [47]. We extend this mathematical interpretation to the physical state of each ncRNA, representing the gene expression as a function throughout the biological sample(i.e., a continuous wave function in time). Our alignment or match function between a 'read' and a ncRNA sequence resembles the inner product or dot product of the vectors (SVs). Although, we are not required to calculate the SVs of the read in the real-time to obtain the alignment, our pre-calculation of SVs combined with the above explained relationship between the SVs and the similarity matrix, would retrieve the same value as if we performed a dot product between SVs, $\{S_A,S_C,S_T,S_G\}$ for ncRNA with the SVs $\{R_A,R_C,R_T,R_G\}$ for the read respectively at the matched region. However, finally to obtain a complex HS, we perform linear addition of all the intermediate HSs obtained for all the mapped reads and the respective ncRNA to obtain the gene expression function i.e., a HS for the ncRNA or simply a DEV. As such, each DEV represents the ncRNA behavior across it's length throughout the sample/cell. DEVs, not only reveal the ncRNAs being processed when visualized, but also lets us apply calculus functions directly on them without the necessity for any pre-processing. Finally, after careful observation of the gene expression curves thus obtained, we were able to map a wavelet function (Ricker Wavelet or Mexican Hat wavelet) with scales 18-38 for the microRNA like ncRNA behavior. To further solidify our notion, we point out the extensive relationship that exists between wavelets, wave-functions, HSs, and Vector Spaces [48]. The Ricker wavelet in time domain is represented using equation below [49] and our overall methodology is picturized in figure 13.

$$r(\tau) = \left(1 - \frac{1}{2}\omega_p^2\tau^2\right)\exp\left(-\frac{1}{4}\omega_p^2\tau^2\right)$$

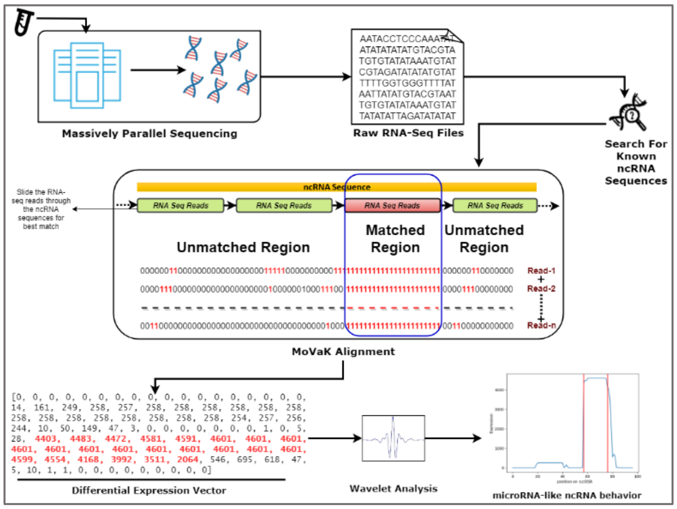

Fig. 13. Workflow For SURFR

#### V. CONCLUSION AND FUTURE WORK

In this paper, we presented a novel, fast, and memory-efficient computational method to identify and analyze the expressions of ndRNAs. Importantly, we demonstrate the significantly improved efficiency of our methodology via comparison to BLAST, and establish our overall accuracy by confirming the identities of our fragment calls in experimentally verified ncRNA fragment databases.

Our method utilizes two principle concepts to identify ndRNAs: SVs for sequence alignment and DEVs for differential expression determination of derived RNAs. We believe, SVs can easily be extended to other use cases of sequence alignment such as alignment with gaps (Insertions/Deletions), or protein alignment etc. Similarly, DEVs can be used as an input for machine learning algorithms such as pattern recognition, clustering, neural networks, or SVMs for mining unknown expression patterns and serious disease associations. Part of our future work will be to demonstrate these use cases. In addition, while we are currently using the existing wavelet, Ricker, for wavelet analysis, we plan to create our own wavelets specific to the behavior of ndRNAs within our DEVs in the near future.

[26] P. McClean, "Basic Local Alignment Search Tool," p. 17.

[27] "Smith-Waterman Algorithm - an overview | ScienceDirect Topics." [Online]. Available: https://www.sciencedirect.com/topics/medicine-and-dentistry/smith-waterman-algorithm. [Accessed: 15-Feb-2019].

[28] S. B. Needleman and C. D. Wunsch, "A general method applicable to the search for similarities in the amino acid sequence of two proteins," Journal of Molecular Biology, vol. 48, no. 3, pp. 443–453, Mar. 1970.

[29] M. Burrows and D. J. Wheeler, "A block-sorting lossless data compression algorithm," 1994.

[30] C. P. C. Gomes, J.-H. Cho, L. Hood, O. L. Franco, R. W. Pereira, and K. Wang, "A Review of Computational Tools in microRNA Discovery," in Front. Genet., 2013.

[38] J. Kinser, Python for Bioinformatics. Jones & Bartlett Publishers, 2010.

[47] R. B. Griffiths, *Consistent quantum theory*. Cambridge ; New York: Cambridge University Press, 2002.

[48] W.-H. Steeb, Hilbert Spaces, Wavelets, Generalised Functions and Modern Quantum Mechanics. Springer Science & Business Media, 1998.

[49] Y. Wang, "The Ricker wavelet and the Lambert W function," *Geophys J Int*, vol. 200, no. 1, pp. 111–115, Jan. 2015, doi: 10.1093/gji/ggu384.
