## Supplementary Information File 3 for "SALTS – <u>S</u>URFR (sncRNA) <u>A</u>nd <u>L</u>AGOOn (lncRNA) <u>T</u>ranscriptomics <u>S</u>uite"

### Supplemental Information File 3. LAGOOn: Algorithm for the identification of lncRNA differential expression and functional prediction

Traditionally, long non-coding RNA computational analysis with respect to anti-sense technology involves four major steps, namely: 1. Performing sequence alignment on raw RNA-seq datasets (two or more) using existing alignment software such as Bowtie or BLAST, 2. Using the aligned-formatted (SAM/BAM) results to obtain the gene expression in terms of reads per million (RPM), 3. Performing statistical analysis to determine the differentially expressed lncRNAs within all the files, 4. Searching the functional-prediction databases based on the differentially expressed lncRNAs. However, the aforementioned steps 1 through 4 requires significant computational expertise either in terms of performing command-line operations or wisely choosing the databases and analytic-parameters with high computational predictability. Also, many of the existing tools are highly generalized in terms of what they perform and involves choosing from many option parameters within the tools. Such generalization of NGS tools account for the requirement of very-high computational resources in terms of memory, processing time, and storing results. For example, a BAM formatted file contains alignment information about all the aligned reads individually plus a lot of other information and, on average, each file after compression is ~128Mb (*BAM File Format*, n.d.). Such general-purpose alignments/tools/results could be an excellent choice for exploratory studies however, pre-defined biological functional analysis requires the domain experts to use the general-purpose software over and over in order to analyze the exponentially growing NGS datasets. Plus, it is absolutely challenging to understand the information from such huge datasets to find/retrieve any patterns which may be significant for biological analyses especially by manually looking at all the aligned reads individually. To address these challenges, we developed a significantly faster, and biologically effective computational method/algorithm in order to analyze RNA-seq datasets (surfr citation). Our algorithm works by interpreting gene expression as a function in four-dimensions(ACTG) on a Hilbert space representing the entire activity of a RNA throughout the sample (one whole RNA-Seq dataset). Such interpretation gives a huge mathematical/computational advantage in order to understand the natural transcriptome behavior using cellular-level calculus, Transcriptomic-Calculus, while still accounting for the dynamism of transcriptomic behavior(including the challenges associated with NGS data generation, organism evolution and, non-significant/significant indels occurring within the transcriptome). Figure 1 shows the difference between the state-of-the-art computational interpretation of RNA-Sequencing data analysis (figure 1.A) versus our understanding of transcriptomic data analysis that works by associating ncRNA behavior to shape of the "Gene Expression Curve" (figure 1.B). Such notion arises from a topological point of view of the RNA activity.

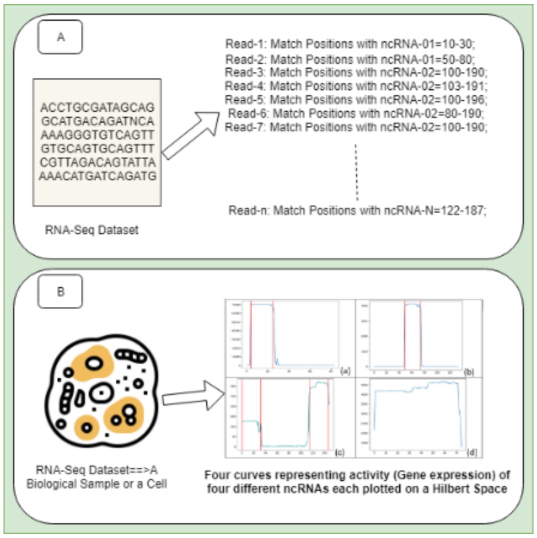


*Figure 1: 1.A) Represents the state-of-the-art interpretation of RNA-Seq data analysis; 1.B) Represents the MoVaK alignment-based Hilbert Space interpretation of the term, "Gene Expression". Whereas, 1.B.a, 1.B.b, 1.B.c represents a "Wavelet" function(s) with scales 18 to 35, which we mapped to microRNA-like ncRNA behavior. However, 1.B.d clearly shows significant difference in terms of ncRNA processing (curve) hence, 1.B.d can be interpreted as a ncRNA that is fully expressed and is not considered microRNA-like behavior.*

Using our theory, not only our algorithm works faster but also our accuracy/precision is compatible with experimental-validation since we are now able to capture the transcriptome behavior using mathematical means as opposed to the traditional understanding of purely string-comparisons/string-alignments (Kasukurthi et al., 2019). However, our methodology was initially engineered specifically towards analyzing small non-coding RNAs and the small RNA fragments that are being derived from them, which significantly differs from the state-of-the-art lncRNA analytic strategies. Similarly, the input requirements and the output interpretations also differ in both the techniques. Hence, we consider this an opportunity to further explain and elaborate on our methodology and show how it can be applied to lncRNA analyses which involves relatively higher computational space and time complexities.

Pre-Processing:

Briefly, our method specific to lncRNA analysis is aimed at determining the expression levels of known lncRNA exons in a given RNA-seq dataset in terms of RPM. In order to do so, we use our own version of seed-and-extend alignment for local alignment of the RNA-Seq reads. Computationally, our alignment with respect to the current study is a modified version of the alignment described in supplementary file one. It works based on two pre-calculated data structures namely, Aho-Corasick Automaton (ACA) for seed search, and Similarity Vectors (SV) for extending the initial seed match. In this section, we explain the significance of these data structures and the steps required to construct them. Our pre-processing stage contains four steps: (1) Data Collection, (2) K-mer separation, (3) ACA construction, and (4) SV construction.

1) Data Collection: Up-to-date annotations of all the human lncRNA exons are obtained from Ensembl. We chose exon sequences rather than the full length lncRNAs since only the exons contain valuable gene expression information which is important for antisense studies. The raw sequence file is then processed using Python 3.7 scripts to extract "ENSE" id for each exon, "ENSG" id for each exon representing the lncRNA it belongs to, and finally the exon sequence (Both ENSE and ENSG are NCBI provided IDs). Then each triplet of ENSE id, ENSG id, and the sequence is associated with another unique id (lagoon id/UID) created by us to be consistent

throughout our software. However, the lagoon id is converted back to the ENSE and ENSG ids when returning the results to the users. Also, any duplicate sequences are separated since it is a redundant task to search for the same sequence.

2) K-mer Separation: It is quite common in RNA-Seq data analysis techniques to filter the reads using short initial seed alignments. This step makes sure that the analysis is only applied to the reads that contain at least a certain exact seed match with one of the RNAs under consideration. In the current study, the seed length we chose is 40nt i.e., any read to be considered for the analysis must contain at least 40nt exact match with one of the lncRNA exons obtained from the previous step. One challenge here is that the 40nt exact match can happen anywhere on the exon so every possible sub-string of length 40 needs to be searched in order not to miss any of the reads. K-mer separation is a step in which we prepare our set of seeds so that the reads can be filtered effectively. Specifically in the current method, to make sure that every possible sub-string of 40nt is searched during the seed-search, we split all the exon sequences into 20nt non-overlapping sub-strings. One of the key differences in terms of pre-processing between our methodology for small ncRNAs (sncRNAs) vs the lncRNAs is that, in sncRNA k-mer separation, we used overlapping sub-strings instead of non-overlapping ones since there are significantly less number of known sncRNAs compared to that of the known lncRNA exons. And the average length of the exons is also substantially big compared to the sncRNAs. So if we were to consider, the total number of overlapping 40-mers for exons would become way too many to be searched efficiently (>10million). Hence, we employed non-overlapping 20-mer separation (<1million) in the current method to trade a large amount of space complexity for a little time complexity. However, every sub-string, either overlapping or non-overlapping needs to be associated with the position of the last character of the sub-string on the exon sequence during k-mer separation.
3) Aho-Corasick automaton construction: AC algorithm is one of the most efficient, widely-used, multiple-pattern (sub-strings) matching algorithms. AC algorithm uses a data structure called "AC automaton" for searching, and is mainly used to search for pre-defined sets of patterns (sub-strings) simultaneously within huge datasets/databases in a very space and time efficient manner. We in our methodology, use AC algorithm to perform the seed search to filter the reads. To do so, ACA needs to be constructed using all the k-mers obtained from the previous step. Our ACA is constructed in such a way that, each search term is indexed using the unique ID of the exon and the search term's last character's position on the lncRNA. Although AC automaton requires significant pre-processing and engineering, once constructed, the automaton can be re-used any number of times. AC algorithm's time complexity is directly proportional to the length of the longest pattern to be searched for, and it's space complexity is directly proportional to the total number of patterns. This is also one of the key reasons to choose non-overlapping 20-mers in the current methodology i.e., in order to keep the size of the created automaton limited (280MB for non-overlapping vs several GB for overlapping mers).
4) Similarity Vector construction: A similarity vector is a data structure we use to hold information regarding the transcriptomic sequences. This data structure is based on the gene-expression information across four dimensions (ACTG) throughout the length of a ncRNA within an entire sample (RNA-seq file). As such, four mandatory similarity vectors are required for each ncRNA sequence to hold information about each of its positions i.e., whether a location contains the alphabet or not. Computationally, a similarity vector is represented using binary arrays as shown in figure 2 that represents an SV for the sequence, "GCCCTCCTGGTGCTTACCACAGGCTGTGTT". Optionally, one vector for each of the other alphabet (within RNA-seq data) can also be considered only if we would like to consider such symbols in our analysis (But in-depth, these symbols are converted back to one of ACTG and

usually only appear because of the limitations within NGS data production). Another noteworthy point to mention is that, SVs for the current study are directly stored in the form of bits rather than arrays of integers to save at least 64 times the space while still acquiring all the benefits of real-time processing.

$S_A$ = [0000000000000001001010000000000] (Match S with A)
$S_C$ = [0111011000000100011010001000000] (Match S with C)
$S_G$ = [1000000001101000000000110010100] (Match S with G)
$S_T$ = [0000100100100110000000000101011] (Match S with T)
$S_N$= [1111111111111111111111111111111] (Match S with N)

*Figure 2: An example of a similarity vector*

**Methodology**

Algorithm:

*Objective*: To identify the expression levels of the known lncRNA exons in a given RNA-Seq dataset.

*Criteria*: Each read must contain 40nt exact match with one of the exons under consideration. And, a read is mapped to an exon only when they share the highest consecutive exact match among all the other (exon,read) pairs i.e., among cross combination of 30,000+ exons vs 10-100million RNA-Seq reads.

*Requirements*: ACA constructed with 20-mer non-overlapping sub-strings, each indexed with the UID (lagoon id) and the sub-string's respective last character position as explained earlier. Plus, SVs for all the exon sequences stored in bit arrays.

*Procedure*:

1. For every given read, ACA-based keyword search is performed to retrieve the following, a) Zero or more exons which has at least 20nt exact match, b) The respective indexed terms, UID and the last character position, c) Finally, the positions on the read where the 20-mers are matched.
2. In the supplemental file one, we presented a relationship that exists among SVs, the diagonals of a similarity matrix and the pair, (last character position of the mer & its respective match location on the read) (also known as position coordinates). We use this relationship to obtain "Alignment Vectors" or "Diagonal elements of the similarity matrix" for each of the exon read pairs from step one. Simply, this step returns multiple binary vectors in which 0 represents a mismatch between the read and the exon at a particular location and 1 represents a match. The pseudocode to do so is shown in fig three.
3. From step 2, a list of exons and associated alignment vectors are returned. Now, the task is to just identify the total number of the most consecutive ones in all the binary lists to obtain the highest consecutive matches among all the exon read pairs. Finally, we consider the exons with the top most highest consecutive 1s only if the number is >=40. Occasionally, more than one exons are returned with the same number of most consecutive ones. But, most of the time, such exons are homologous.
4. For each exon obtained from step 3, increment a global counter representing that a read is mapped to it.
5. Repeat steps 1 through 4 for all the reads in a RNA-Seq file to obtain a list of exons and their respective gene expression in terms of number of reads mapped.
6. Finally, the read counts are converted to reads per million.

```
Given the read 'R', SV {A, C, T, G, N}, length of read 'n', length of ncRNA 'm', list of position coordinates, (position on ncRNA, position on Read) (ai, bi)=[(a1,b1),(a2,b2),(a3,b3)…]
    //Initialize an empty set data structure (holds only unique values)
    SET E[];
    //Initialize a 1d matrix of length m with 0s.
    Alignment=[0] of length 'm'
    //repeat for every tuple of position coordinates
    for each (ai, bi) {
    //if the difference of ai, bi in E then skip
    if (ai-bi) in E:
    continue;
    else:
    //add the value a-b to the set
    add (ai-bi) to E;
    j= n-(bi-ai)-1;
    k= m-(ai-bi)-1;
    start1= min(j+1,m);
    stop1= max(1,j-n+2);
    start2= max(1,k-m+2);
    stop2= min(k+1,n);
    // step = -1 gives output in reverse order
    //we use python's "arange" function instead of a for loop here
    //arange(1,4)=[1,2,3,4]      whereas      arange(1,4,-1)=[4,3,2,1]
    //x= all the x coordinates of the diagonal elements.
    //y= all the y coordinates of the diagonal elements.
    x= for(start1-1; stop1-2; step= -1);
    y= for(start2-1, stop2-2; step= -1);
    //Initialize another 1d matrix to hold diagonal element values
    Diagonal=array (length m);
    for (int val=0; val < length(x); val++){
    //Fill in the diagonal elements using an SV and positions (xj,yj)
    Diagonal [x[ val ] ]= SV [R [ y [ val ] ] ] [x [ val ] ];
    }
    //Perform 1d matrix addition
    Alignment= Alignment + Diagonal;
```

*Figure 3: Pseudocode For Aligning A Read With A Known Sequence Using SVs*

**Discussion & Real-Time Usage Of Our Algorithm:**

The steps mentioned above requires only one iteration through the entire RNA-Seq file. The time complexity is bound by three major factors 1. The ACA search complexity, 2. Identification of highest consecutive ones in binary numbers, and 3. The rate of hardware IO capability. However, since ACA time complexity is linear, and highest consecutive ones in binary number can be achieved using bit operations and is also linear (Plus, something to note here is, it takes no real-time comparisons or computations at all to extend the initial seed match using SVs), one of the crucial time consuming factor in the above algorithm is the rate at which the hardware supports the IO operations. For example, on solid state drives, our algorithm performed at least 40% faster compared to that of traditional hard drives. Due to the low memory and time requirements, our algorithm is capable of real-time speedups consuming only several minutes to satisfy all the requirements necessary for lncRNA analyses. Finally, as a proof-of-concept we developed a user-friendly web application that uses the above procedure to automate the lncRNA analysis including real-time differential expression analysis, and we made it available for free of cost at salts.soc.southalabama.edu/lagoon_home. The algorithm development, data structure creation, and all the other computations are performed using Python 3.7 programming language and, the results are stored, retrieved, & compared using mongoDB document store (The Most Popular Database for Modern Apps, n.d.).

Two of the main differences between our methodology for sncRNA analysis and lncRNA analysis are k-mer separation, and the biological implication of the read alignments (RPM vs The Gene Expression Curve). To be specific, we've shown how time and space complexities in our methodology can be traded accordingly based on the requirement, for example, k-mer separation step has been modified strategically to include significantly low seed count (space complexity) while still achieving linear time-complexity. And, when it comes to the specific biological significance of the aligned reads at particular locations on the transcriptome, we've achieved it by identifying microRNAs & like RNAs using wavelet functions on four-dimensional vector space by extending our alignment vectors into Differential Expression Vectors (See supplementary file one), which again is a Hilbert space interpretation of the term "transcriptome expression" in a given sample.

Finally, we have also included lncRNA functional predictions in our tool to fulfill the state-of-the-art requirements. Specifically, we chose five characteristics of lncRNAs that are highly examined namely, lncRNA exons as sponge, lncRNAs as host, lncRNA overlapping Genes, lncRNA interactions with enhancers, and lncRNA micro-protein analysis using Ribo-Seq data analysis. All the components in the toolkit are described in figure four. Each of these components, and how they are integrated into the tool are briefly described below.

*lncRNA exons as sponge:* lncRNAs frequently function as miRNA sponges that directly basepair with and effectively inactivate mature miRNAs. Due to this we have identified all miRNA complementarities of 10 base pairs of greater occurring within each of the lncRNAs contained within our database. The sponge dataset has been generated using miRBase mature miRNAs (Griffiths-Jones et al.).

*lncRNAs as host*: Numerous lncRNAs have been shown to encode sncRNAs (e.g. miRNAs and snoRNAs) in their exonic sequences and sncRNA expression to rely on excision from the host lncRNA's transcript. As such, we have identified full length ncRNAs contained with lncRNA exons. The host dataset has been generated using Ensembl Biomart (Yates et al.).

*lncRNA interactions with enhancers*: Several lncRNAs have been reported to function through regulating the accessibility of transcriptional enhancers overlapping their genomic loci. All overlaps between a given lncRNA and annotated enhancer are therefore described. The dataset for this feature has been generated using GeneHancer (Fishilevich et al.).

*lncRNA overlapping genes*: In addition to lncRNA exonic sequences serving as sncRNA hosts, many sncRNAs are processed from lncRNA introns. Furthermore, many lncRNAs serve as naturally occurring antisense silencers of genes located on the strand opposite to themselves. For both of these reasons, as well as a few other potential regulatory relationships, all genes overlapping a lncRNA locus on either the positive or negative strand are indicated. The overlapping genes dataset has been generated using Ensembl(Yates et al.).

*lncRNA micro-protein analysis using Ribo-Seq*: For the users to perform micro-protein analysis in-addition to the differential gene expression analysis of the lncRNAs, we have extended our algorithm to Ribo-Seq data analysis as well. The criteria we used for Ribo-Seq analysis is 30nt exact match and extended to find the best match. As such, we used the same exon dataset as earlier for the Ribo-Seq functionality.

Finally, all the datasets generated above, the results from real-time, user-uploaded datasets (Both Ribo-Seq and lncSeq) are integrated using the unique IDs mentioned earlier (ENSG, ENSE, lagoon ids) over mondoDB engine for efficient querying.

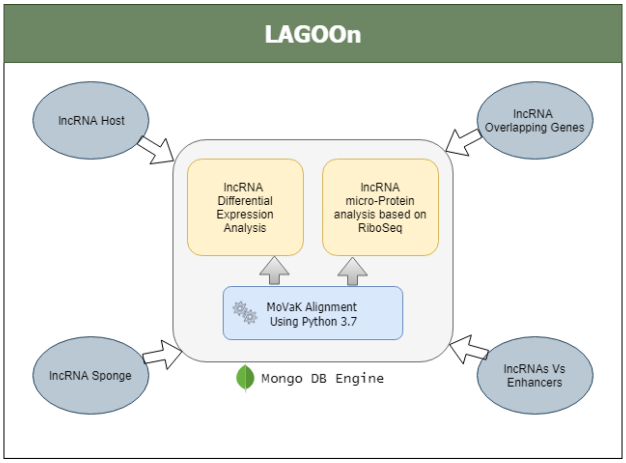


*Figure 4: Overview Of The Components In LAGOOn*
